# Integrated analysis of different non-coding features across the *Sox2* locus implicates a diencephalic enhancer in adult brain expression

**DOI:** 10.1101/680082

**Authors:** D.A. Carter

## Abstract

SOX2 is a prominent member of the SOX family of transcription factors that has many different functional roles. This pleiotropy is made possible by multiple regulatory mechanisms that direct appropriate spatial and temporal patterns of expression, and therefore action. The current study concerns the mechanisms that determine *Sox2* gene expression in the adult mammalian brain, where SOX2 protein is absent in general, but is selectively and abundantly expressed in a majority of neurons within a ventral diencephalic brain structure, the suprachiasmatic nucleus (SCN). In this study, a comparative bioinformatic and biochemical analysis of different adult rat brain regions was conducted in order to identify SCN-selective (immaturity-related) regulatory mechanisms. The approach incorporated an integrated analysis of *Sox2* enhancers, CTCF binding sites, and also expression of the *Sox2*-overlapping, long non-coding (lnc)RNA, *Sox2ot*. Initial experiments revealed brain region-specific *Sox2ot* expression (including region-specific novel transcripts), indicating a significant diversity of *Sox2ot* expression across the adult brain. However, the pattern and abundance of *Sox2ot* expression in the SCN, relative to selected control areas of the brain, did not indicate an overt relationship to *Sox2* gene expression. Furthermore, although multiple individual *Sox2ot* exon sequences were shown to overlap annotated *Sox2* gene enhancers at different sites across the *Sox2* locus, again there was no indication of a SCN-specific functional correlation. Further integration with an analysis of selectively-active CTCF sites within the *Sox2* locus directed attention to one site with both a prominent peak of activity in immature brain, and proximity to a functionally-characterized, ventral diencephalic, *Sox2* enhancer termed U6 (upstream enhancer 6). *Ex vivo* analysis of the U6-associated CTCF site revealed SCN-selective CTCF binding, and these sequences were both localized within a known (brain region-selective) super-enhancer. Bioinformatic analysis of the U6 enhancer sequence revealed an abundance of consensus sites for the SCN-selective transcription factor LHX1, and over-expression of this factor enhanced the activity of cloned U6 sequence in transfected cells. However, despite this compelling evidence for a molecular mechanism that underlies adult brain expression of SOX2, further analysis of LHX1-SOX2 co-expression in the SCN confounded this view, indicating the presence of other concurrent mechanisms in the different cell populations of the SCN.

## INTRODUCTION

SOX2 (sex determining region Y-related high-mobility group (HMG) box 2) is a transcription factor with established developmental roles both in the early embryo (Avilion et al, 2003), and also in later processes including neurogenesis in the developing brain (Cavallaro et al, 2008; Favaro et al, 2013; Sarkar & Hochedlinger, 2013). In accordance with a role in early neuronal development, expression levels of the *Sox2* gene generally fall as neurons reach final maturation, and, *ipso facto*, SOX2 has a limited role in the adult brain. However, our work, and that of others (see Hoefflin & Carter, 2014; Cheng et al, 2019), has identified that *Sox2* expression is maintained in specific adult neurons; in particular, there is abundant SOX2 within one cluster of neurons in the adult rodent brain, the suprachiasmatic nucleus (SCN) of the ventral diencephalon (Hoefflin & Carter, 2014; Cheng et al, 2019). In fact, adult SCN neurons represent an interesting example of apparent neuronal immaturity where, in addition to the expression of developmental factors such as SOX2, and the cytoskeleton regulator doublecortin (Geoghegan & Carter, 2009), these cells also escape postnatal silencing of paternal *Ube3a* alleles, found in the majority of adult neurons (Jones et al, 2016). The unusual expression pattern of SOX2 in the SCN has recently been afforded greater significance, firstly through the finding of a similar pattern of SOX2 expression in the SCN of adult human brain (Pellegrino et al, 2018), but, moreover, through functional analysis in mice which showed that SCN SOX2 is required for robust circadian time-keeping, the cardinal function of this brain region in mammals (Cheng et al, 2019).

Further research is now required to identify the molecular mechanisms that control SCN *Sox2* expression, i.e. the mechanisms that permit both the escape from common neuronal gene silencing (Hoefflin & Carter, 2014), and also the role in adult circadian function (Cheng et al, 2019). Such an analysis has to be conducted with reference to an extensive body of known mechanisms that regulate the *Sox2* gene, which involve both conventional gene enhancers, and also other novel, non-coding sequences. The *Sox2* gene is located within a gene desert region (Ovcharenko et al, 2005), and multiple, distinct enhancer sequences have been identified over a substantial 200kb chromosomal region around the *Sox2* coding sequence (see Okamoto et al, 2015). However, the specific individual roles of the 27 identified neural *Sox2* enhancers (see Okamoto et al, 2015) are not fully identified (Sugahara et al, 2018; Tomioka et al, 2002; Uchikawa et al, 2003; Zhou et al, 2014). Similarly, the contribution of other modes of gene regulation is also undefined. These include a ‘super-enhancer’ organization that is known to regulate some cell-specific expression of *Sox2* (Li et al, 2014), but remains partially defined. Secondly, there is a regulatory RNA component that involves the *Sox2*-associated, long non-coding RNA (lncRNA), *Sox2ot* (Sox2-overlapping transcript; Amaral et al, 2009) which spans the *Sox2* locus, but is also ill-defined with respect to cell-specific transcript structure, and expression. Functionally, *Sox2o*t, like other lncRNAs, may act either *in cis* (Clark & Blackshaw, 2014; Joung et al, 2017) or *in trans* (Briggs et al, 2015) to regulate *Sox2* expression, and currently there is some evidence of both types of mechanism (Askarian-Amiri et al, 2014; Messemaker at al, 2018). The recently identified *cis*-level mechanism in mouse embryonic stem cells (Messemaker at al, 2018) involves an interaction of *Sox2ot* with a *Sox2* enhancer, in accordance with a widespread recognition of enhancer-lncRNA interactions (Vucicevic et al, 2015; Cajigas et al, 2018). Therefore, in view of this large body of data, there is clearly a need to conduct an integrated analysis of the multiple potential *Sox2* regulatory sequences that are likely to be involved in permitting adult neuronal expression of this gene.

The current study has directly investigated *Sox2ot* transcript structure and expression in the SCN, and related this data to *Sox2* enhancer organization. Transcriptomic analysis has previously indicated that *Sox2ot*, unlike *Sox2*, is highly expressed in the adult brain; it is one of the top 50 most highly expressed lncRNAs in adult mouse brain (Kadakkuzha et al, 2015), where expression has been likened to ‘housekeeping’ (Liu et al, 2016). However, the distribution pattern of *Sox2ot* in the adult brain is undefined. Previous studies have demonstrated the presence of multiple different *Sox2ot* transcripts, each composed of differentially spliced exons that derive from >500kb of the *Sox2* locus, and it is clear that other variants remain to be discovered (Amaral et al, 2009; Shahryari et al, 2015). As brain *Sox2ot* variants are likely to exhibit cell-type specificity of expression (Raj & Blencowe, 2015), it will be interesting to determine any association between SCN-selective exon usage, and the location of individual *Sox2* enhancers, which have been shown to have brain-region specific activities (Okamoto et al, 2015). The current analysis of *Sox2ot* and *Sox2* enhancers has also been integrated with an analysis of CTCF (CC<u>CTC</u>-binding <u>F</u>actor) activity at the *Sox2* locus because recent studies have shown that one of the multiple activities of CTCF is to control differential expression of neuronal genes (Hirayama et al, 2012). The current studies were conducted on rat brain because this species has been the focus of our previous work (Geoghegan & Carter, 2009; Hoefflin & Carter, 2014), and the size of the rat brain facilitates selective brain tissue sampling of the SCN for the biochemical analysis required in this study.

## METHODS

### Animals

Adult male rats (Sprague-Dawley, postnatal day 50) were used in accordance with the UK Animals (Scientific Procedures) 1986 Act of Parliament, and the study was also approved by the Cardiff University ethical review committee. Animal health was monitored by a veterinarian, and rats were maintained in standard laboratory conditions (14:10 light:dark cycle, lights on: 05.00h) with *ad libitum* access to rodent food and drinking water. For both RT-PCR (3 rats/sample) and Chromatin immunoprecipitation (ChIP) analysis (4 rats/sample), rats were killed by a Schedule 1 method at 17.00h, and punched samples of SCN and cerebral cortex (parietal area; COR) tissue were obtained using a rat brain matrix (RBMA-300C, World Precision Instruments Inc., Sarasota, FL, USA) and blunt 15G needles (Holter et al, 2001). In addition, the olfactory bulb (OB) was sampled *in toto* for use in RT-PCR analysis. Samples for RT-PCR were stored at −80°C prior to RNA extraction. Samples for ChIP analysis were used immediately for chromatin preparation. Unless otherwise stated, analyses were conducted on three independent biological replicates. For immunohistochemical analysis, animals were anaesthetized with sodium pentobarbitone (150 mg/kg, i.p), and perfused via the ascending aorta with phosphate buffered saline (PBS), followed by 4% paraformaldehyde in 0.1M phosphate buffer (PFA). Whole brains were dissected, post-fixed in PFA (overnight, 4°C), cryoprotected (20% sucrose in 0.1M phosphate buffer, overnight, 4°C), and then stored briefly at –80°C prior to sectioning. Cellular expression of proteins was confirmed in multiple brain sections from 3 adult rats.

### RT-PCR analysis

RNA was extracted from rat brain samples using Trizol (Invitrogen, Thermo Fisher Scientific, Waltham, MA, USA) and DNaseI-purified (Promega, Madison, WI, USA). cDNA was generated using the Superscript II protocol (Life Technologies, Thermo Fisher Scientific) using an Oligo (dT) primer. PCR analysis was conducted using standard procedures; the Q5 Hot-Start High-Fidelity DNA polymerase (NEB, Ipswich, MA, USA), and 1% agarose gel electrophoresis. Oligonucleotides used for PCR amplifications are listed in Table S1. Amplified products were visualized with reference to DNA ladders (either: Hyperladder I, Bioline, London, UK, or 1kb ladder, Promega), using GeneSnap (Syngene, Frederick, MD, USA). PCR products were then cut from gels, purified (Qiaex II gel extraction kit, Qiagen, Hilden, Germany) and ligated into pGEM-T (Promega protocol). Ligations were transformed into JM109 cells (Promega), and colonies were selected for plasmid purification (PureYield protocol, Promega). PCR products were then sequenced (Eurofins Genomics, Ebersberg, Germany).

### Analysis of Sox2 enhancer activity

Transcriptional enhancer activity of the rat ‘U6’ sequence was tested by transient transfection in a reporter construct expressed in hippocampal HT-22 cells (Li et al, 1997) using methods previously employed in our laboratory (El Kasti et al, 2012). The U6 reporter construct was assembled by ligating a 663bp rat U6 sequence into the HindIII and NheI sites of the pGL4.23 vector (Promega) upsteam of the minimal promoter in this vector. The U6 sequence was amplified from rat genomic DNA using the primers U6TFF1 and U6TFR3 (Table S1), and sequence-verified (Eurofins Genomics). HT-22 Cells (1×10^5^/well of a 24-well plate) were grown in DMEM with 10% Fetal Bovine Serum, and 1x antibiotic/antimycotic (Invitrogen) at 37°C and 5% CO_2_, and co-transfected (TransFast protocol, Promega) with the test constructs (PureYield plasmid, Promega protocol), and pRL-TK (5ng) as a control. In some experiments, cells were co-transfected with a (mouse) LHX1 expression construct (Origene Technologies Inc., Rockville, MD, USA) in which Myc-DDK-tagged-LHX1 is expressed from pCMV6. Following transfection, cells were then maintained for 24h prior to lysis and luciferase assay according to the manufacturers protocol (Dual-luciferase reporter assay system, Promega). Relative luminescence values were measured on a Luminometer (Model TD-20/20, Turner Biosystems, Sunnyvale, CA, USA). Each transfection was replicated 6-fold (3 replicates, in 2 separate experiments). The data was analysed by first normalizing against the individual pRL-TK values, and then calculating fold-difference compared to the activity of ‘empty’ pGL4.23 vector. In other experiments, cells were maintained for 24h after transfection, and then fixed in 4% formaldehyde (Sigma) and processed for immunocytochemical detection of LHX1 protein as described below.

**Fig.1.**
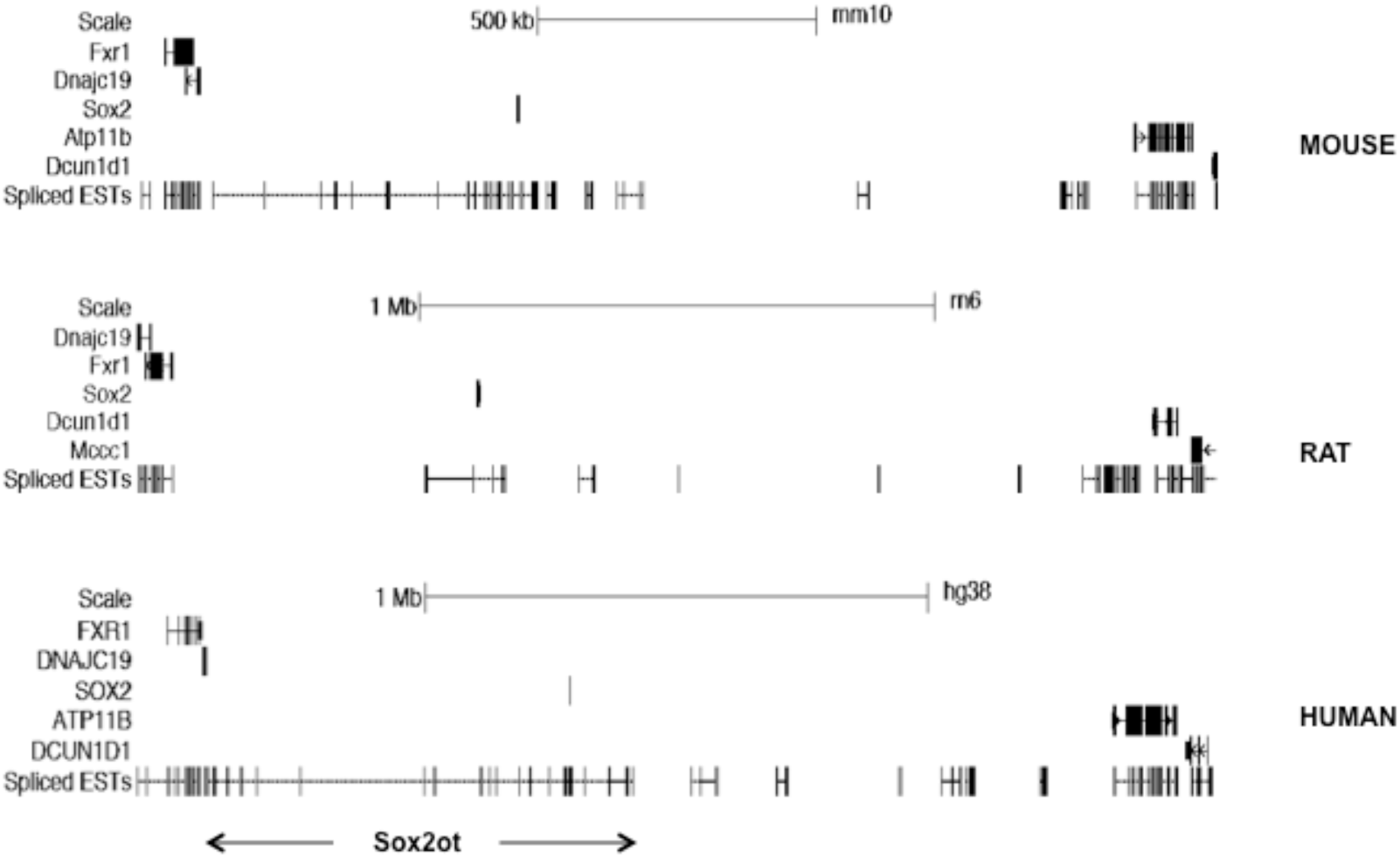
*Sox2* locus structure is conserved across mammalian genomes. UCSC genome browser images showing that *Sox2* is located within a >1Mb ‘gene desert’, but is overlapped by transcribed sequences (Spliced ESTs in ‘dense’ format) that form the incompletely annotated *Sox2ot* lncRNA.

**Fig.2.**
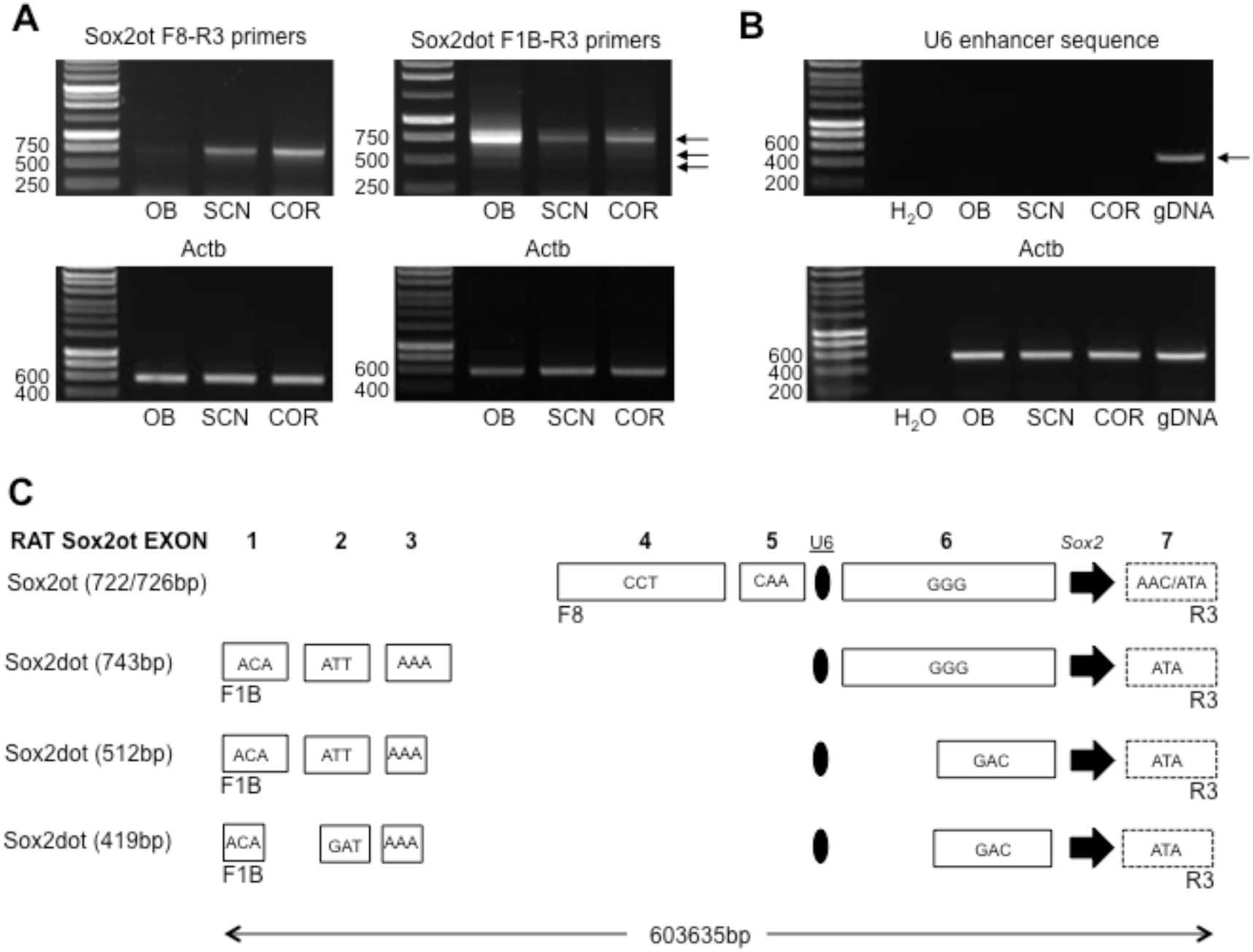
*Sox2ot* transcripts are formed from exons that are widely distributed across the *Sox2* locus, and differentially incorporated in different rat brain regions. **A.** Representative images of agarose gel electrophoresis analysis of PCR-amplified *Sox2ot* (left, F8-R3 primers) and *Sox2dot* (right, F1B-R3) transcripts in olfactory bulb (OB), suprachiasmatic nucleus (SCN) and cortex (COR). Note the low levels of *Sox2ot* product in OB, but high relative levels of *Sox2dot* product. RT-PCR analysis of *Actb* transcripts are shown for comparison. Numbers on the left of gels are sizes in base pairs. B. Representative images of agarose gel electrophoresis analysis of PCR-amplified U6 enhancer sequence. Note the absence of product in OB, SCN and COR cDNA compared with the definitive 522bp product amplified from genomic DNA (gDNA). **C.** Schematic representation of rat *Sox2ot/dot* exon structure. Alternative PCR product sizes are shown on the left (see text), and each exon is depicted by a rectangle that includes the first 3 bases of the exon sequence. The 3’-truncated exon 7 (see text) is dashed. The *Sox2* gene and U6 enhancer are represented by an arrowhead and oval respectively. Note that exon sizes have an indication of scale, but the distribution across the 604kb of rat genome is not drawn to scale.

### Chromatin Immunoprecipitation analysis (ChIP) analysis

ChIP and associated PCR analysis was conducted as described in our previous study (Davies et al, 2011) with minor modifications. Each brain sample for chromatin preparation was dissected as described above, and composed of either SCN punches from 4 rats, or an equivalent volume of cortex. The ChIP-IT Express kit (Active Motif, Carlsbad, CA, USA) was used as described (Davies et al, 2011), but here a glycine-addition step was incorporated, as indicated in the kit protocol. Chromatin was sheared to a 300-600bp fragment preparation, in clear contrast to a preparation of rat genomic DNA (Sambrook et al, 2001; Fig.2A) using a closed system ultrasonic disruptor (10 cycles of 30s ON/30s OFF, 4°C; Bioruptor Pico, Diagenode sa, Liege, Belgium). ChIP assays were then conducted using antisera to CTCF (61311, Active Motif), histone H3K27ac (39133, Active Motif), LHX1 (C6, Santa Cruz Biotechnology, Santa Cruz, CA, USA), and IgG as a control (ChIP-IT Control Kit, Active Motif). Use of the CTCF antibody in ChIP analysis of brain chromatin was initially verified using a known CTCF target sequence in the *Igf2* gene (Ling et al, 2006). Semi-quantitative analysis was conducted using standard PCR amplification (Q5 Hot-Start High-Fidelity DNA polymerase; NEB, Ipswich, MA, USA, and agarose (2%) gel electrophoresis. PCR primers were designed around target sequences (Table S1), and gel images were obtained using GeneSnap software (Syngene, Frederick, MD, USA). PCR band sizes were verified using a molecular mass DNA ladders (Hyperladder, Bioline Ltd., London, UK; Quick-Load Purple 100bp ladder, NEB) and band intensity was measured using ImageJ (Schneider et al, 2012). Initial validation in *Igf2* was conducted only in single replicate samples, but subsequent analysis of the *Sox2* locus was conducted in triplicate, using three independent biological samples.

### Immunohistochemical and immunocytochemical analysis

For rat brain analysis, coronal sections of brain containing the SCN were cut on a cryostat (Leica CM1900; Leica Imaging Solutions Ltd., Cambridge, UK), and mounted on glass slides (SuperFrost Plus, VWR International Ltd., East Grinstead, West Sussex, UK). Slides were briefly dried, stored at −80 °C, and immune-histochemical analysis conducted as described (Hoefflin and Carter, 2014). For immunocytochemical analysis of transfected HT-22 cells, fixation (4% formaldehyde, 15 min.), and detection (Hoefflin and Carter, 2014 protocol) was conducted *in situ* on the culture surface of 12-well Costar CellBIND® plates (Corning, Kennebunk, ME, USA).

The SOX2 primary antibody (39823, Active Motif, Carlsbad, CA, USA) has been validated for immunohistochemical analysis in our previous study (Hoefflin and Carter, 2014). The LHX1 primary antibody (C6, Santa Cruz Biotechnology) was used by this laboratory for the first time here, and was validated by demonstrating a previously characterized, cell-specific, nuclear expression in the adult SCN (Vandunk et al, 2011). The antisera to both betaIII-tubulin (G7121, Promega) and nuclear histone (H3K27Ac, 39133, Active Motif) which were used to visualize cell populations in the immunocytochemical analysis, were validated by characteristic cytoplasmic, and nuclear localization respectively. In addition, all antisera were subject to simple control validation by demonstrating both excitation light-specific-detection, and absence of detection when the primary antibody was omitted. The secondary antisera used were: Cy3-conjugated donkey anti-mouse IgG (Jackson Immunoresearch Laboratories Inc., West Grove, PA, USA) and Alexa Fluor 488-conjugated goat anti-rabbit IgG, (Molecular Probes Inc, Eugene, OR, USA). Following final washing, both brain sections and fixed cells were mounted using Vectashield with DAPI (Vector Laboratories, Burlingame, CA, USA), and stored at 4°C. Brains and fixed cells were viewed with a fluorescence microscope (Leica DM-LB, Leica), and images were captured with a Leica DFC-300FX digital camera linked to Leica QWin software (V3). Representative images were assembled in Photoshop (CS2, Adobe Systems Inc., San Jose, CA, USA).

### Bioinformatics and statistical analysis

Genomic and transcript analysis was conducted on the UCSC and Ensembl Genome Browsers (genome.ucsc.edu; ensembl.org), using different genome assemblies, where required, for specific annotation, including ENCODE (www.encodeproject.org). Analysis of cloned DNA sequences was conducted using BLAST (blast.ncbi.nlm.nih.gov), and Clustal Omega (ebi.ac.uk/Tools/ msa/clustalo). Either MatInspector (Genomatix Software Gmbh, Munich, Germany) or Lasagna (biogrid-lasagna.engr.uconn.edu), as available, were used to identify transcription factor consensus binding sites. VISTA enhancer sequences were obtained from the VISTA enhancer browser (enhancer.lbl.gov). Gene expression patterns were viewed at both the Allen Brain Atlas (mouse.brain-map.org) and GTEx (V6; gtexportal.org). PCR primers were designed using Primer-BLAST (ncbi.nlm.nih.gov/tools/primer-blast). Statistical analysis was conducted with IBM SPSS Statistics version 20 (IBM, New York, USA) using different tests as described in Results, and applying a significance cut-off of p<0.05.

## RESULTS

### Sox2 is located in a conserved gene desert region of rat chromosome 2

In order to define the *cis*-regulatory region around the rat *Sox2* gene, current genome browser views of the *Sox2* locus in rat, mouse and human were compared (Fig.1). This *in silico* comparison shows that, in each species, the *Sox2* gene is located within a similar ‘gene desert’ region of >1Mb, and is overlapped by transcribed sequences (ESTs) that form the incompletely annotated *Sox2ot* lncRNA.

### Sox2ot is highly expressed in rat brain

In initial experiments, the expression of *Sox2ot* transcripts was analysed in the rat brain, comparing expression in samples extracted from olfactory bulb (OB), suprachiasmatic nucleus (SCN) and cortex (COR) (Fig.2). Accurate sampling of the SCN was validated by showing SCN-selective exclusion of rat *Grin1* exon 4 in a PCR analysis (Fig. S1), as demonstrated in a previous study (Partridge and Carter, 2017). In subsequent comparisons of *Sox2ot* expression in the three brain regions, a complex pattern of expression, distinct for different regions, was observed.

For *de novo* amplification of *Sox2ot* sequences from rat brain, PCR primers were designed with reference to rat EST sequences that have homology to mouse *Sox2ot* (CB556907.1 and DY317483.1), Allen Brain Atlas *in situ* hybridization probe sequences (Experiments: 79906731 and 79906733), the sequence of a known mouse cDNA (BC057611) and *Sox2ot* sequences reported in previous work by Amaral and colleagues (Amaral et al, 2009). An end-point PCR approach using agarose gel electrophoresis was selected because of the need to identify, and discriminate between anticipated *Sox2ot/dot* splice variants (Shahryari et al, 2015). Using this approach, two broad classes of rat Sox2ot transcripts were detected, similar to those identified in mouse and human: (i) transcripts overlapping and proximal to *Sox2*, termed *Sox2ot* transcripts, and (ii) transcripts overlapping, but incorporating (5’) distal exons, termed *Sox2dot* (see Amaral et al, 2009; Shahryari et al, 2015). Cloning and sequencing of these PCR products revealed both differential exon incorporation, and exon length (Fig.2; Supplemental data, S2).

### Sox2ot transcript expression

Using the primer pair SoxotF8 & SoxotR3, *Sox2ot* transcripts were readily amplified from rat brain, but with distinct regional differences in RNA levels. Thus, whereas levels of the ∼700bp F8R3 amplicon in SCN and COR were similar (COR levels are 114.3±13.2% of SCN levels, both corrected for *Actb* amplicon levels p=0.366, Independent samples t test, n=3 brain samples), levels were markedly lower in olfactory bulb (Fig.2A). This result was consistently observed in PCR analysis of each of 3 independent brain samples but the OB product was insufficient for accurate quantitation in some samples. Cloning and sequencing of the ∼700bp PCR product from each brain region revealed two very similar, but distinct sequences. The 4-exon, rat *Sox2ot* cDNAs (Fig.2A&C; Supplemental data, S2) are similar to previously described human 4-exon Sox2ot transcripts (see Shahryari et al, 2015), and appear to fully exclude an additional intermediate exon (between rat exons 6 & 7, Fig.2C.) compared with known human (and mouse) 5-exon Sox2ot transcripts (see Shahryari et al, 2015). It is recognized that the amplified sequence of rat exon 7 (ATA/AAC start) does not represent a full-length exon sequence; additional PCR amplifications with distal 3’ primers (data not shown) indicate that this exon is similar in size to the final exon of the mouse cDNA (BC057611), which extends to >2kb.

Surprisingly, the rat *Sox2ot* transcripts identified here exhibited subtle, region-specific variation, producing the two variants indicated above: (i) cortex transcripts lacked the initial 4 bases of exon 7 that were present in (ii) SCN and OB transcripts (Supplemental data S2). This variation is not found in known human and mouse sequences, and also contrasts with a predicted rat transcript (LOC103691527, variant X5); therefore the cortex F8R3 Sox2ot product is distinct and appears to be representative of a novel transcript (Genbank Accession: MH204884). While it is possible that this 4bp variation could represent a somatic variation in cellular sub-populations, it is more likely a tandem alternative splicing event (Szafranski & Kramer, 2015) given the presence of two closely spaced ‘AG’ sequences at the start of rat exon 7 (see Supplemental data, S2). With respect to potential sequence function *in trans*, the 4bp variation may be functionally relevant because loss of the 4bp significantly modifies potential peptide sequence encoded by *Sox2ot/Sox2dot* variants (see below; Supplemental data S3). In contrast, this variation would have minimal apparent effect on selected RNA binding protein (RBP) sites (Supplemental data S4). Most obviously, however, the markedly lower levels of F8R3-spanning *Sox2ot* transcripts in OB relative to SCN and COR would appear to be significant with respect to any potential *in trans* actions of this specific lncRNA sequence.

### Sox2dot transcript expression

Using the primer pair SoxotF1B & SoxotR3, 5 exon, rat *Sox2dot* cDNAs were amplified (Fig.1A&C; Supplemental data, S2) that are similar to previously described human *Sox2dot* transcripts, but lack some intermediate exons that be representative of human/tissue-specific transcripts (see Shahryari et al, 2015), For *Sox2dot* transcripts, a more extensive sequence variation compared with rat brain *Sox2ot* transcripts was observed; three distinct PCR products were obtained of 743, 512 and 419bp, which incorporate variations in exon sequence, and fully exclude rat exons 4 & 5 (Fig.1A&C, Supplemental data, S2). However, rat Sox2dot transcripts did not exhibit the ATAG deletion from the start of exon 7 (Supplemental data, S2). Of note, the truncated exon 3 in two of the *Sox2dot* variants utilizes a non-canonical splice donor site (Supplemental data S2). The 3 sequenced rat brain *Soxdot* variants are similar to, but distinct from a predicted rat transcript (LOC103691527, variant X9), and therefore novel. Interestingly, the most abundant (743bp) Sox2dot transcript was present at markedly higher levels in OB, compared with SCN (23.3±4.7% of OB level, n=3) and COR (31.0±3.6%, n=3); samples (Fig.1A). Relative quantitative analysis of this result, corrected for equivalent levels of *Actb* amplicon, revealed a significantly lower level of the 743bp product in SCN and COR relative to the OB level (p<0.05, ANOVA followed by Duncan’s test; F=(2,8) 37.596). With respect to potential sequence function i*n trans*, this reduction may be functionally relevant because it would significantly lower levels of a potential peptide sequence encoded by *Sox2dot* (Supplemental data S3). In addition it is possible that a lower abundance of cellular RNA sequence could affect free cellular RBP activity (Supplemental data, S4).

Overall, these results indicate that the rat has similar *Sox2ot* transcripts to other species, but also exhibits a number of exonic sequence variations that may be rat-specific. Comparative analysis across different brain regions has not indicated both quantitative and qualitative differences between SCN, OB and COR samples, but there was no major distinction between the (abundant Sox2-expressing) SCN, and (control) COR samples. However, the presence of high (rat brain *Sox2ot* mRNA levels are approximately 10% of Actb mRNA levels, Sutherland & Carter, unpublished), and regionally-variable *Sox2ot* expression prompted an analysis of the relationship of the *Sox2ot* transcript structure to enhancer sequences associated with the *Sox2* locus.

### Relationship between brain *Sox2ot* expression, and *Sox2* enhancers

Analysis of the relationship between *Sox2ot* expression in rat brain and the location of known *Sox2* enhancers is feasible because the published functional enhancers are highly conserved across chicken and mammalian genomes (Okamoto et al, 2015). In the latter study, the authors provided detailed evidence of sequence conservation across chicken, human and mouse genomes. Here, the presumed conservation in rat was confirmed by direct sequence comparison with rat genomic sequence. For example, the U6 *Sox2* enhancer identified by Okamoto and colleagues exhibits extensive DNA sequence identity across human, mouse and rat sequences (Supplemental data S5). Consequently, given the mouse and human sequence information provided by Okamoto et al (2015), together with the conservation between rat and mouse *Sox2ot/dot* exons (see above), it has been straightforward to map expressed rat *Sox2ot/dot* exons to predicted enhancer locations in both the mouse and human genome browsers (genome.ucsc.edu), where ENCODE annotation is available (www.encodeproject.org).

This bioinformatic analysis has shown that, for *Sox2ot* exons, there is often a direct correspondence between expressed exons and a sub-set of the annotated enhancer sequences (Fig.3). Thus, *Sox2ot/dot* exon sequences 4 and 5 overlap U9/10, and exon 6 sequences overlap the N3[C4] predicted neurosensory *Sox2* enhancers, whereas exon 7 terminates proximal to, but just upstream of SC2[C23] (Fig.3). For the *Sox2dot* distal exons, the same correspondence cannot be mapped because these exons are outside the region mapped in the Okamoto *et al* study (2015). However, by mapping these expressed exons to ENCODE annotation of the mouse and human genomes (Fig.3, Supplemental data S6 and Fig. S7), it can be seen that *Sox2dot* exons 1, 2 and 3 are either within, or proximal to predicted/putative enhancer sequences. Thus, exon 1 is within a conserved forebrain-specific Vista enhancer annotated on the human genome (192; enhancer.lbl.gov, Supplemental data S6), and both exons 1 and 2 are within ENCODE-annotated, brain-specific, ChIP-Seq-derived peaks of H3K27ac and H3K4me1 marks (Fig.3C; Supplemental Fig. S7). Exon 3 is somewhat distinct, being located upstream of a potential enhancer sequence with a H3K4me1 peak in brain (Fig. 3C).

**Fig.3.**
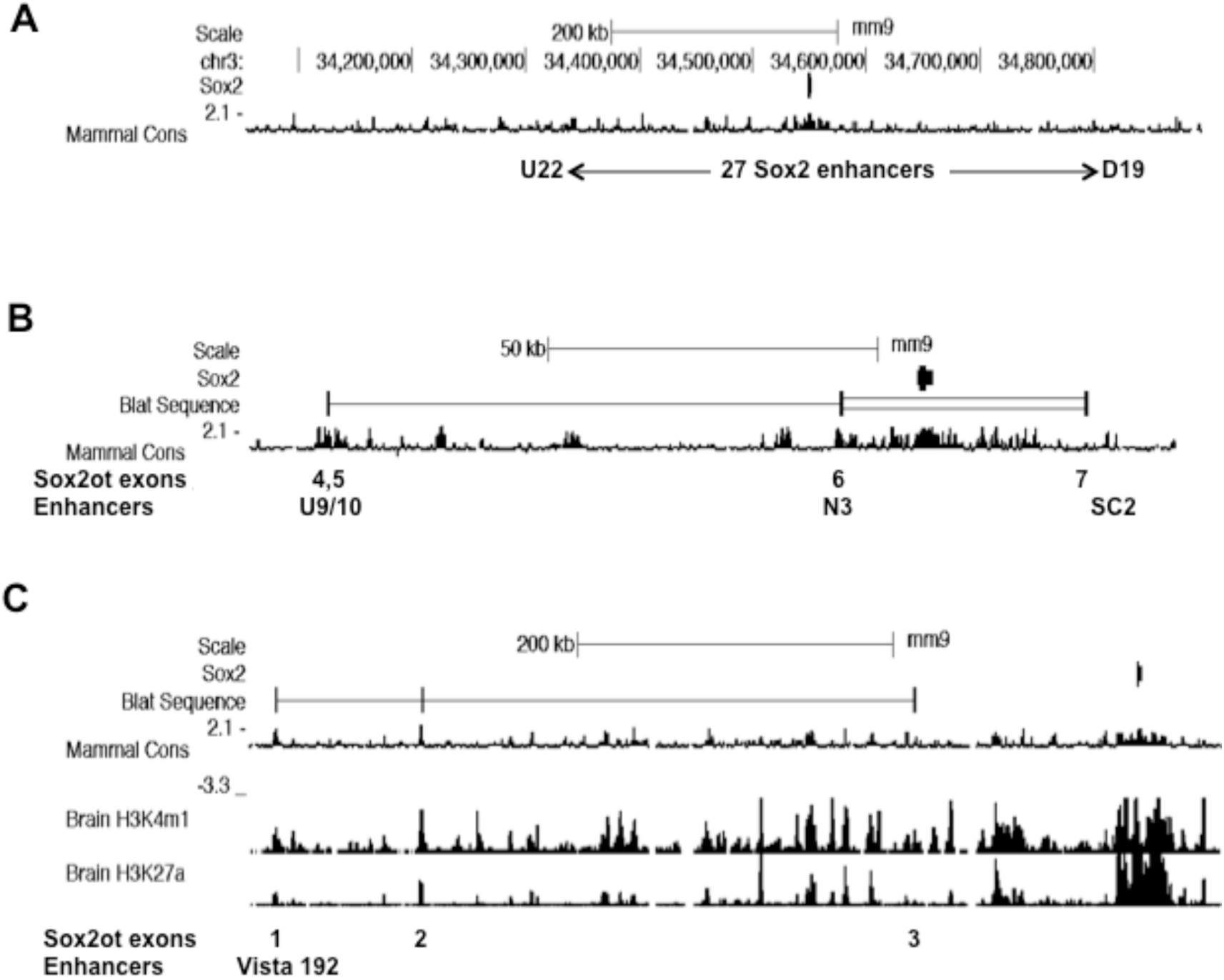
A subset of *Sox2ot* exons overlap Sox2 enhancer sequences. UCSC genome browser images (mouse, mm9) showing the location of *Soxot* exons with respect to conserved *Sox2* enhancer sequences. **A.** Image showing the genomic region mapped by Okamoto et al (2015) that contains a total of 27 neural *Sox2* enhancers, from U (upstream) 22 to D (downstream) 19. **B.** Image showing locations of *Sox2ot* exons (BLAT input = exons 4,5,6,7) relative to conserved *Sox2* enhancer sequences as indicated by conserved blocks of sequence (phyloP). **C**. Image showing the locations of *Sox2ot* exons (BLAT sequence input = exons 1,2,3) relative to conserved sequence (phyloP), and ChIP-Seq analysis of brain H3K4me1 and H3K27ac association (ENCODE/LICR data).

The current demonstration of *Sox2ot/dot* transcription from *Sox2* enhancer sequence is supportive of a role for this lncRNA in the regulation of *Sox2* expression, but immediately raises the question of why the majority of the highly conserved 27 neuro-sensory *Sox2* enhancers (see Okamoto et al, 2015) are not associated with *Sox2ot* transcripts. Consequently, a search was conducted for annotated transcripts (ESTs etc) across the *Sox2* locus (mouse, mm10) in order to identify potential transcripts that may be missing from currently annotated *Sox2ot* sequences. This analysis revealed very limited evidence of additional (known) enhancer-associated transcription, either sense or antisense; just two antisense lincRNAs (Gm29135-201 and Gm43207-201) partially overlap the U17 and U4 enhancers, respectively (Supplemental data S8). Notwithstanding a requirement for deep sequencing of this transcriptome from relevant tissues, the current combination of RT-PCR and bioinformatic analysis therefore indicates two distinct classes of *Sox2* enhancer, those transcribed within *Sox2ot*, and those not so transcribed. However, it must be recognized that the (albeit extensive) identification of *Sox2* enhancers by Okamoto and colleagues (2015) is incomplete. In order to provide an indication of the full number of *Sox2* enhancers, a bioinformatic search was conducted over the entire span of the Sox2 locus (i.e. from the 5’-flanking coding gene: *Dnajc19*, position 33,980,110 on mm9, to the 3’-flanking coding gene, *Atp11b, position* 35,653,306 on mouse genome [mm9]), a total genomic distance of 1.67Mbp. This search was conducted using mm9 because the UCSC-based, ENCODE data can provide an indication of putative enhancers through the coincidence of peaks of H3K4me1 and H3K27ac binding. This *ab initio* analysis revealed a total of 64 such regions across the locus, where 8 were coincident with genome-annotated *Sox2ot* exons, and a further 6 were also coincident with Sox2ot exons, but additionally coincident with a H3K4Me3 peak, indicating a possible *Sox2ot* promoter (rather than enhancer) sequence. This preliminary analysis on the mouse genome indicates an abundance of uncharacterized enhancers across the *Sox2* locus, and supports the above conclusion that a substantial number, albeit a minority, of *Sox2* enhancers are included within *Sox2ot* transcripts. Further experimental collation of both *Sox2* enhancers and *Sox2ot* exons is required to fully define the relationship between these two entities.

### Relationship between Sox2 enhancers and CTCF activity

Given the absence of any overt, SCN-selective, relationship between adult *Sox2ot* transcript expression and annotated *Sox2* enhancers, a third parameter (CTCF binding) was introduced for analysis in order to highlight potential sequences with tissue-selective properties. This approach was based on recent evidence of CTCF binding at lncRNA-associated enhancers (see Ntini & Marsico, 2019), and the recognition that tissue-specific CTCF activity can mediate differential expression of neuronal genes (Hirayama et al, 2012, Prickett et al, 2013). The current analysis involved a survey of ChIP-seq-derived CTCF DNA binding on the ENCODE database, focused on major ChIP-Seq CTCF peaks observed in either ES cells or embryonic brain (Fig.3). This revealed (at the resolution selected in Fig.4A) ten peaks of maximal amplitude, of which six exhibited similarly high activity in peripheral tissues, including liver. Notably, however, four other peaks exhibited brain/ES cell specificity, and one peak (labelled 4, Fig.4A) was of particular interest because the activity was high in both ES cells and embryonic brain, but very low in adult brain, indicating a site that could be selectively active in immature neurons. This site was of further interest because, in contrast to the other selected CTCF peaks, it is proximal to one of the annotated *Sox2* enhancers (U6, Fig. 4B, see Ntini & Marsico, 2019). Moreover, the U6 enhancer is a highly conserved (mouse vs. human, 81%, Supplemental data S5) and functionally verified, ventral diencephalic enhancer sequence (Okamoto et al, 2015). Given this specific functional relevance to the current study of ventral hypothalamic (SCN) *Sox2* expression, an *ex vivo* analysis of CTCF activity at the U6-associated site was conducted in samples of adult rat brain.

**Fig.4.**
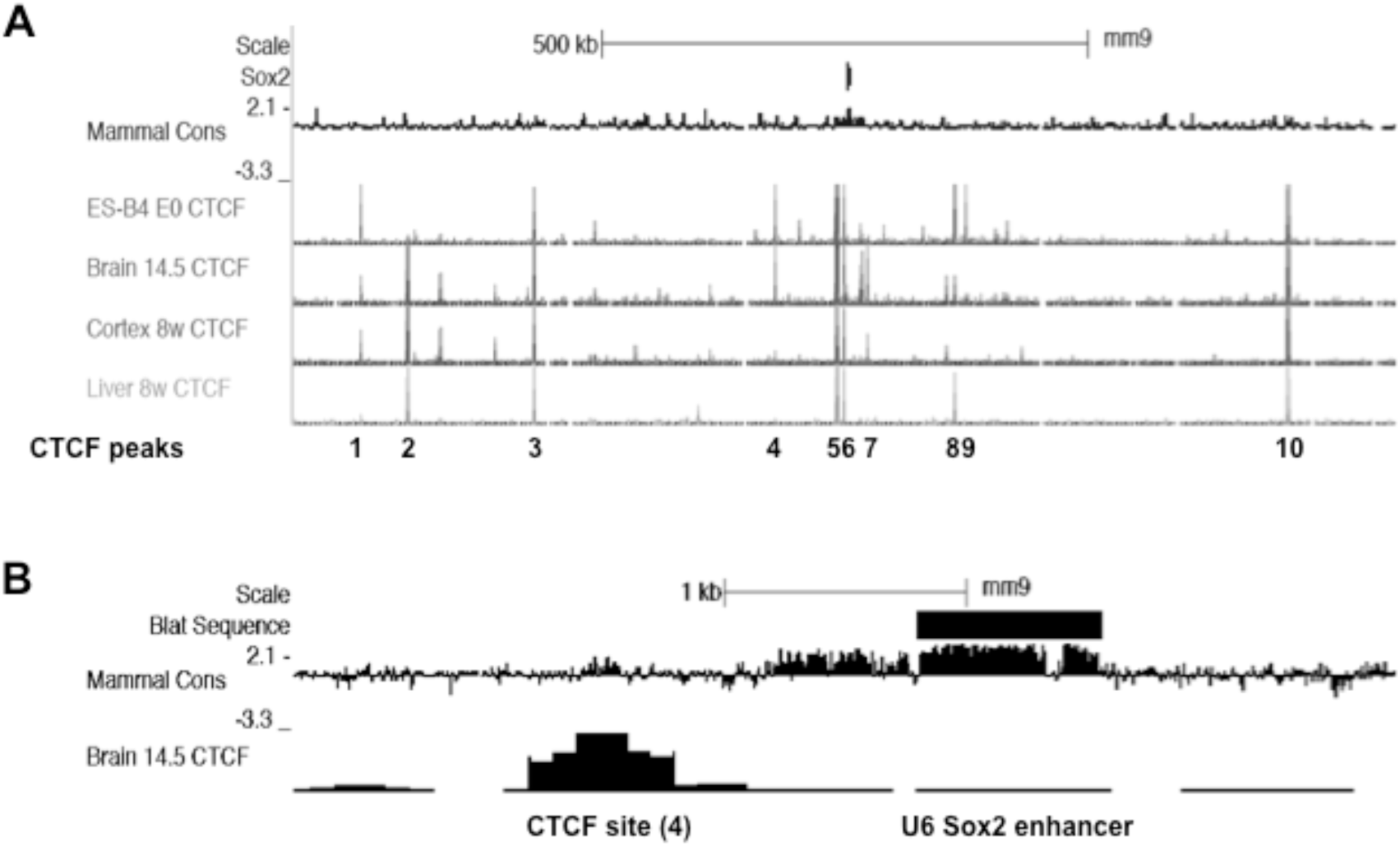
CTCF ChIP-seq analysis across the Sox2 locus reveals tissue/developmental-specific peaks of association. UCSC genome browser images (mouse, mm9) showing CTCF ChIP-Seq peaks across the Sox2 locus. **A**. Among 10 major peaks of CTCF association, peak 4 is notable for high levels in ES cells and embryonic brain (Brain [day] 14.5) but low levels in adult brain (cortex 8w) and liver. **B.** Detail of (A) showing the proximity of CTCF site 4 with the U6 *Sox2* enhancer. Note the conservation of mammalian sequence at the core of the CTCF site, and across the U6 enhancer region.

### ChIP analysis of CTCF activity in adult rat brain

In an initial part of this analysis, further evidence of potential CTCF activity at the U6-associated site was obtained by conducting an *in silico* consensus transcription factor binding site analysis of the U6-associated, ChIP-Seq-derived CTCF sequence. This analysis revealed that consensus CTCF sites are present in both mouse and rat U6-associated sequence (Supplemental data S9). Also, to provide a control sequence for experimental ChIP analysis (see below), additional sequence analysis of the major, *Sox2*-proximal, ChIP-Seq-derived CTCF site (labelled 5, Fig.4A) also revealed conserved CTCF consensus sites in rat (Supplemental data S9). In order to provide experimental (*ex vivo*) evidence of CTCF activity in the adult rat brain, a ChIP analysis was then conducted using chromatin extracted from adult brain (Fig.5). In preparation for this analysis, an appropriate level of chromatin shearing in the brain samples was verified by agarose gel electrophoresis (Fig.5A), and a commercial CTCF antibody was validated for ChIP analysis using a published CTCF target sequence in the *Igf2* gene (Ling et al, 2006; Fig.5B). Following these control analyses, antibody-specific immunoprecipitation for both the distal, U6-associated CTCF site, and the major proximal CTCF site was then demonstrated (Fig.5C). Subsequently, it was shown that whereas the proximal CTCF site exhibited a similar level of association in both cortex and SCN, only SCN samples exhibited CTCF association at the distal, U6-associated site (Fig.5D). To provide additional context for this result, an additional ChIP analysis of a major H3K27ac-associated site (part of the Sox2 super-enhancer [Li et al, 2014], located downstream of Sox2; Supplemental data S10), demonstrated antibody-specific chromatin association, but, in this case, no difference in the levels of immunoprecipitated sequence between brain cortex and SCN (Fig.5D). Quantitative analysis of this result revealed no significant difference between SCN and cortex (SCN amplicon level expressed as a percentage of the cortex level: SCN= 93.2±4.1%; p=0.167(p>0.05), Independent samples t test; n=3 independent samples). Taken together, these results indicate relative tissue-specificity for CTCF association at the U6-associated site within SCN neurons, and therefore the current study focused on this site in further analyses.

**Fig.5.**
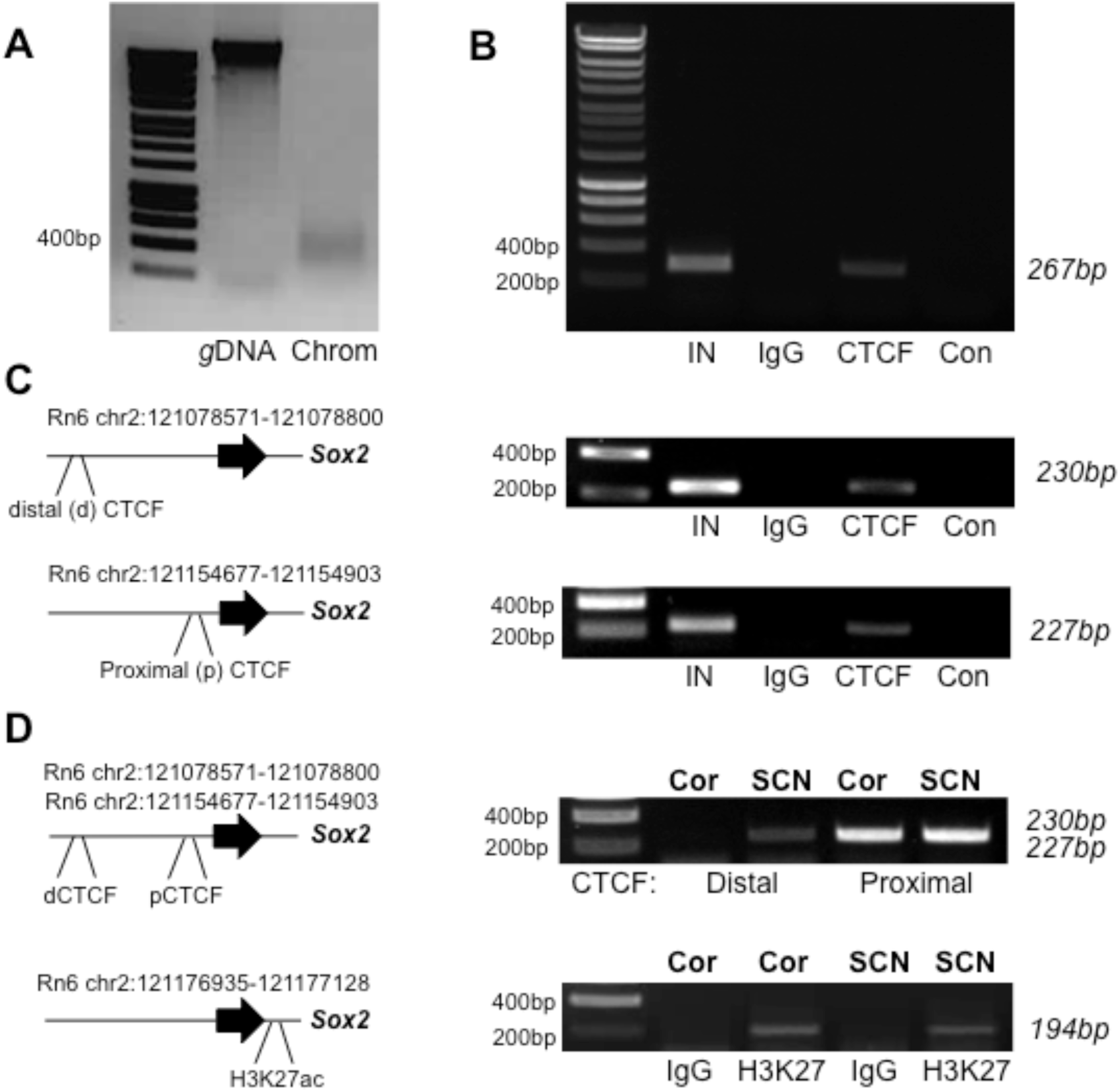
CTCF is differentially associated at target sites within the *Sox2* locus in SCN and cortex. Representative images of agarose gel electrophoresis analysis of ChIP experiments performed on rat brain samples. **A**. Inverted image of gel electrophoresis of rat brain chromatin (Chrom, 2.5µg) used in ChIP analyses, compared with a sample of rat genomic DNA (gDNA). Note that the chromatin sample is present as a smear around the 400bp band. **B.** Validation of the CTCF antibody for ChIP analysis using SCN-derived chromatin, and PCR primers amplifying a 267bp amplicon that includes a known CTCF target site in the *Igf2* locus. Note the CTCF antisera-specific amplicon. **C.** Validation of the ChIP analysis of both distal (upper) and proximal (lower) CTCF sites associated with the *Sox2* locus. Schematic diagrams show the approximate locations of the sites relative to the *Sox2* gene (denoted by arrow). Note the CTCF-specific amplicons for each site**. D.** Comparison of CTCF (upper) and histone H3K27ac (lower) association with the *Sox2* locus in cortex and SCN. For CTCF, note the SCN-specific product for the distal site, and similar product abundance in SCN and Cor for the proximal site. For H3K27ac, note the similar product abundance in SCN and Cor. Abbreviations: Con, water PCR; Cor, brain cortex; CTCF, CTCF antibody; H3K27, histone H3K27ac antibody; IgG, antibody control IgG; IN, input chromatin PCR; Prox, proximal CTCF site relative to Sox2 gene; SCN, suprachiasmatic nucleus. Note that in each case, Input chromatin (IN) on the gel is at a final dilution of 1/80 relative to the ChIP-derived chromatin sample. Number values are sizes in base pairs (bp).

### The U6 enhancer is associated with a novel brain super-enhancer

Scrutiny of the U6 enhancer region within ENCODE annotation of the mouse genome revealed an association with a larger cluster of ChIP-seq-derived H3K27ac marks that are indicative of super-enhancer status (Fig.6A). This presence of a super-enhancer is supported by additional ENCODE annotation of the human genome, most prominently by the presence of a major cluster of H3K4me1 marks around the U6 sequence that are cell-type-specific, and not associated with gene promoter (H3K4me3) marks (Fig. 6B). A search of the dbSUPER database of human super-enhancer activity (Hnisz et al, 2013) also supported this inference, revealing overlap with an annotated human super-enhancer sequence (Fig.7A). The latter finding is intriguing because the Hnisz et al. (2013) study also provided evidence of tissue-specific, super-enhancer status, being positive for human cingulate gyrus, but not for equivalent samples of hippocampus or caudate nucleus (Hnisz et al, 2013). This large, super-enhancer region is also of interest in being proximal to the start of the *Sox2ot* transcript illustrated in Fig.6A.

**Fig.6.**
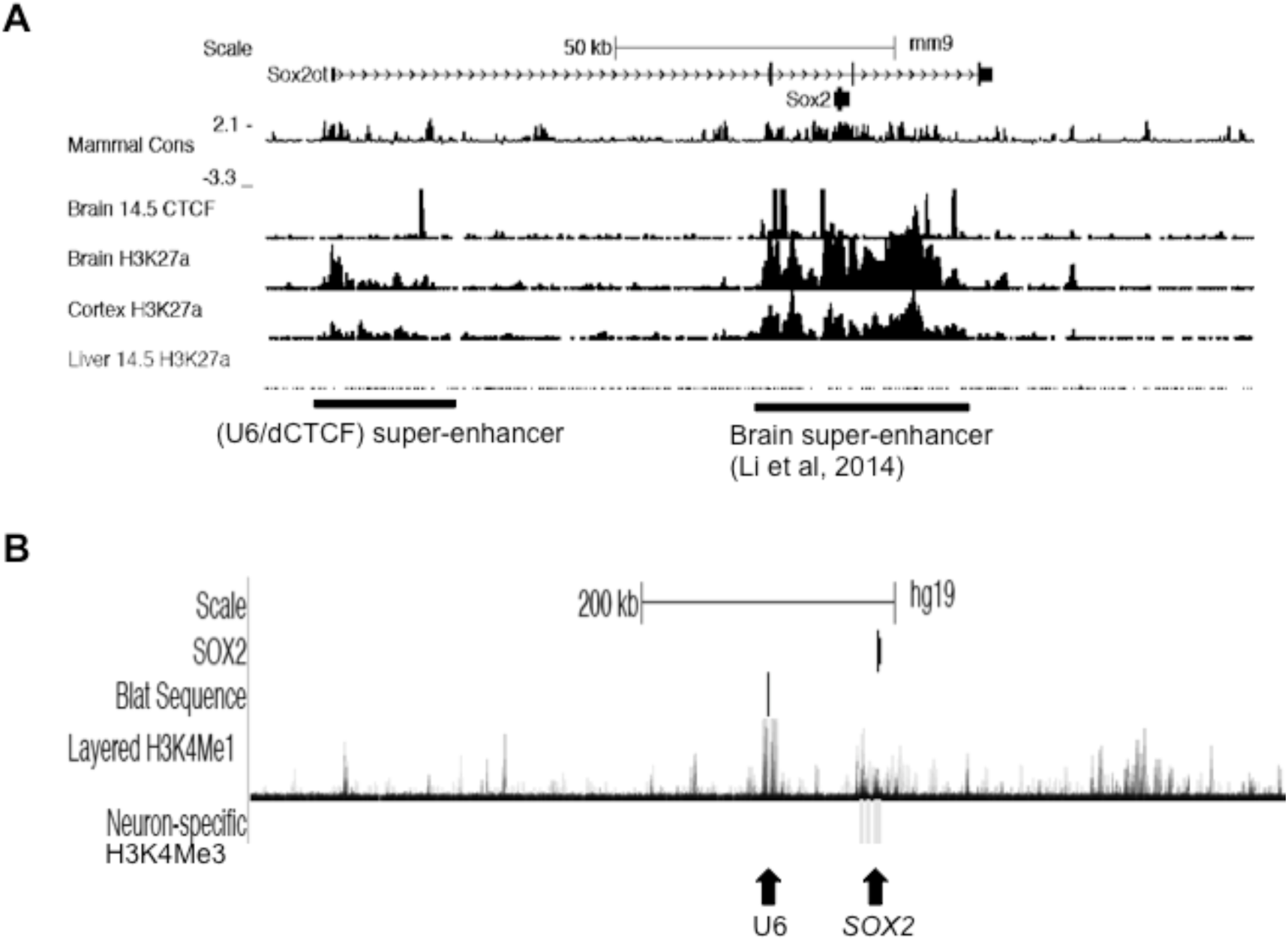
Identification of a super-enhancer region associated with the U6 *Sox2* enhancer. UCSC genome browser images (mouse, mm9; human, hg19) showing evidence of super-enhancer features. **A.** Clusters of H3K27ac associations surrounding the U6 enhancer region. U6 is proximal to the isolated peak of embryonic brain CTCF association. Note the major brain super-enhancer region (Li et al, 2014) proximal to the *Sox2* gene. **B.** A prominent cluster of H3K4me1 marks proximal to the U6 region (U6 identified by Blat sequence). Colours indicate different cell-types (ENCODE); note that the prominent H3K4me1 marks (ENCODE) are cell-type specific (HUVEC and K562). In contrast, neuronal H3K4me3 marks (Brain Histone H3K4me3 ChIP-Seq from Univ. Mass. Medical School (Akbarian/Weng), denoted by ‘neuron-specific’), considered an active mark for promoters, are restricted to the region proximal to the *Sox2* coding sequences.

### Functional analysis of the U6 enhancer region

The evidence obtained above is indicative of a role for the U6 enhancer in regulating adult brain, and possibly SCN-specific, expression of *Sox2*. The current RT-PCR analysis of adult rat brain *Sox2ot* (see above) indicates that U6 sequences are not included within *Sox2ot/dot* transcripts, because the U6 region is flanked by the currently used *Sox2ot/dot* primer sequences. However, in order to confirm an absence of U6-associated sequence within *Sox2ot*, or other, transcripts, further RT-PCR analyses were conducted using U6-intrinsic primer sequences (Table S1). An absence of amplified product in either OB, SCN or COR samples indicates either an absence, or very low transcription from this sequence (Fig. 2B). This result is consistent with an absence of EST sequence at this locus (genome.ucsc.edu), and also with the concept that U6 encompasses a ‘conventional’ enhancer sequence that would be anticipated to bind specific transcription factors. An analysis of consensus transcription factor binding sites within the U6 sequence was conducted in comparison with all other annotated diencephalic enhancers (Okamoto et al, 2015; N3[C4], D1, N2 C[8], N2[C9], and N3[C2]). This analysis revealed an abundance of three different consensus sites in the U6 sequence, but only one of these (LHX3) was 100% conserved across the rat, mouse and human genomes (Supplemental data S11 & S5). In contrast, there was no enrichment of LHX3 sites in the other diencephalic enhancer sequences (Supplemental data S11). This selective enrichment of consensus LHX3 sites has potential significance because the LHX core binding motif is identical across the different LHX factors (Berger et al, 2008), and one member of this family, LHX1 (LIM homeobox 1), has a specific role in SCN development and function (Bedont et al, 2014; 2017), and in the adult brain is selectively expressed in the SCN (Allen Brain Atlas, Experiment 79591731; mouse.brain-map.org; Furuyama et al, 1996).

Although the U6 enhancer region is highly conserved (Supplemental data S5) and has been functionally verified as an enhancer in embryonic electroporation experiments (Okamoto et al, 2015), additional experiments were conducted to confirm enhancer activity of rat U6 sequence with respect to LHX1 interaction. After demonstrating low relative levels of *Lhx1* expression in non-transfected HT-22 cells (Fig. 7B), it was shown that rat U6 sequence mediated a significant increase in transcriptional activity from a minimal promoter in pGL4.23, but only in the presence of enhanced levels of LHX1 (Fig. 7C & Supplemental Fig. S12).

**Fig.7.**
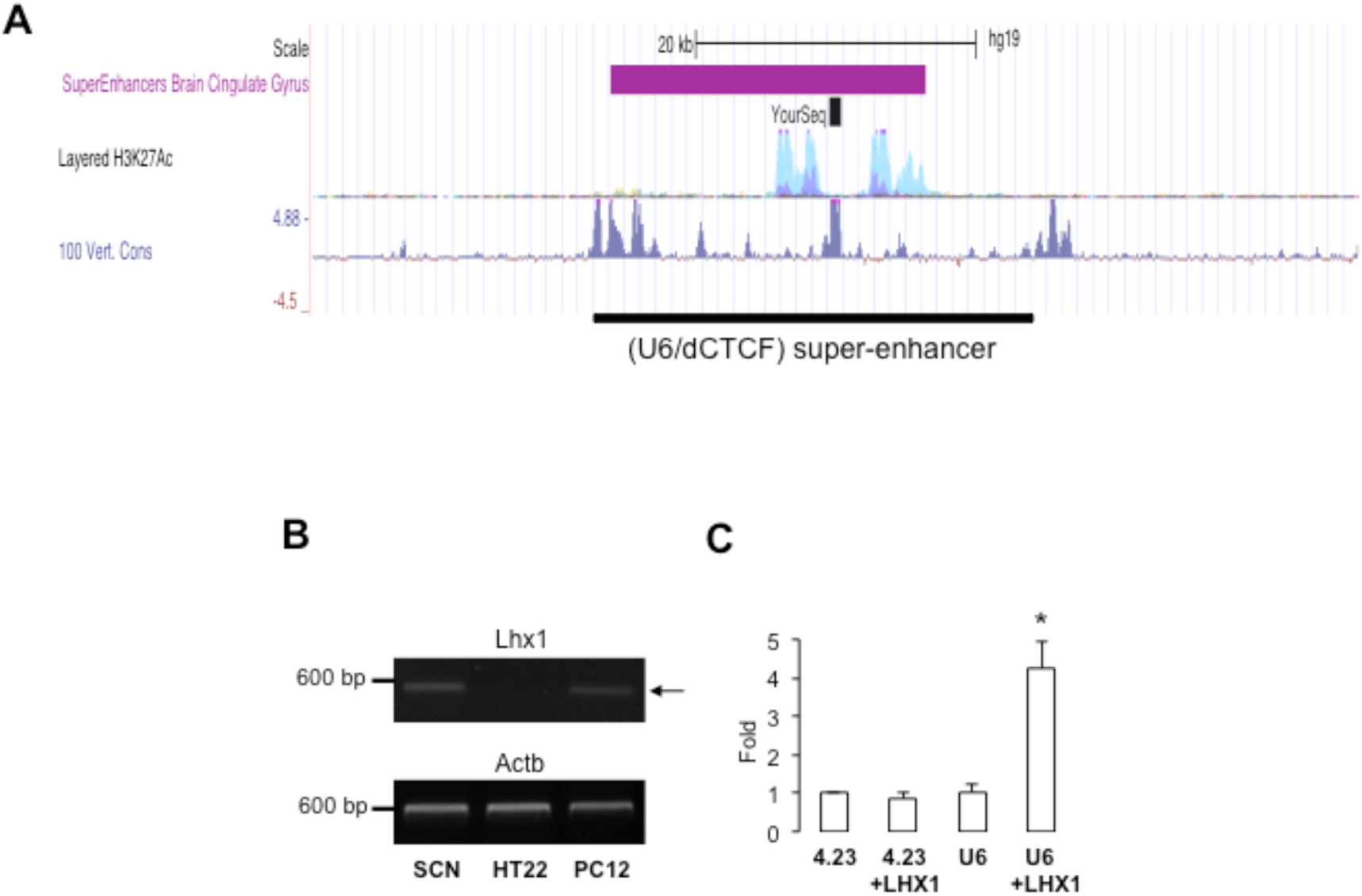
Functional analysis of the U6 *Sox2* enhancer. **A.** The wider genomic region that contains U6 (the U6 sequence is denoted by ‘Your Seq’ [BLAT search]) has been identified as a super-enhancer for human cingulate gyrus (Hnisz et al, 2013). **B.** RT-PCR analysis of *Lhx1* mRNA expression in neuronal cell lines indicates low relative expression in HT-22 cells. Representative images of agarose gel electrophoresis analysis of PCR-amplified *Lhx1* and *Actb* in suprachiasmatic nucleus (SCN), and HT-22 and PC12 cells. Numbers on the left of gels are sizes in base pairs. **C.** The rat U6 enhancer sequence mediates an enhanced transcriptional response from pGL4.23 when co-transfected with an LHX1 expression plasmid in HT-22 cells. Histogram showing fold-change in relative luciferase activity in different conditions compared with the empty pGL4.23 plasmid (n=6/group; * = p<0.05 compared with all other groups; ANOVA and Duncan’s test, F=(3,20) 16.357.

### LHX1 expression in the rat SCN, relative to SOX2 expression

The evidence obtained above, and associated literature, is indicative of a functional relationship between LHX1 (through U6 sequence) and SOX2 in the SCN of the rat. In order to investigate the relative expression of these two factors, an immunohistochemical analysis was conducted in the adult rat brain where SOX2 is highly, and unusually expressed in the SCN (Hoefflin & Carter, 2014; Fig.8). This analysis confirmed that LHX1 is also extensively expressed in the adult rat SCN (Fig.8), a highly selective pattern of expression given the absence of immunoreactivity both in the cortex (Supplemental Fig. S13) and all other brain regions viewed in these ‘SCN-region’ coronal brain sections. Clear co-localization of LHX1 and SOX2 was observed in many neurons (Fig.8E-H). However, SOX2 is much more extensively expressed across the SCN; LHX1 immunoreactivity is primarily confined to a ventromedial sub-region of the SCN as previously demonstrated for the mouse (Vandunk et al, 2011). Consequently, there are numerous SOX2+ neurons in the dorsal SCN where LHX1 is absent or minimally expressed (Fig. 8H), and therefore LHX1 does not appear to be obligate for the maintenance of SOX2 expression in the adult SCN.

**Fig.8.**
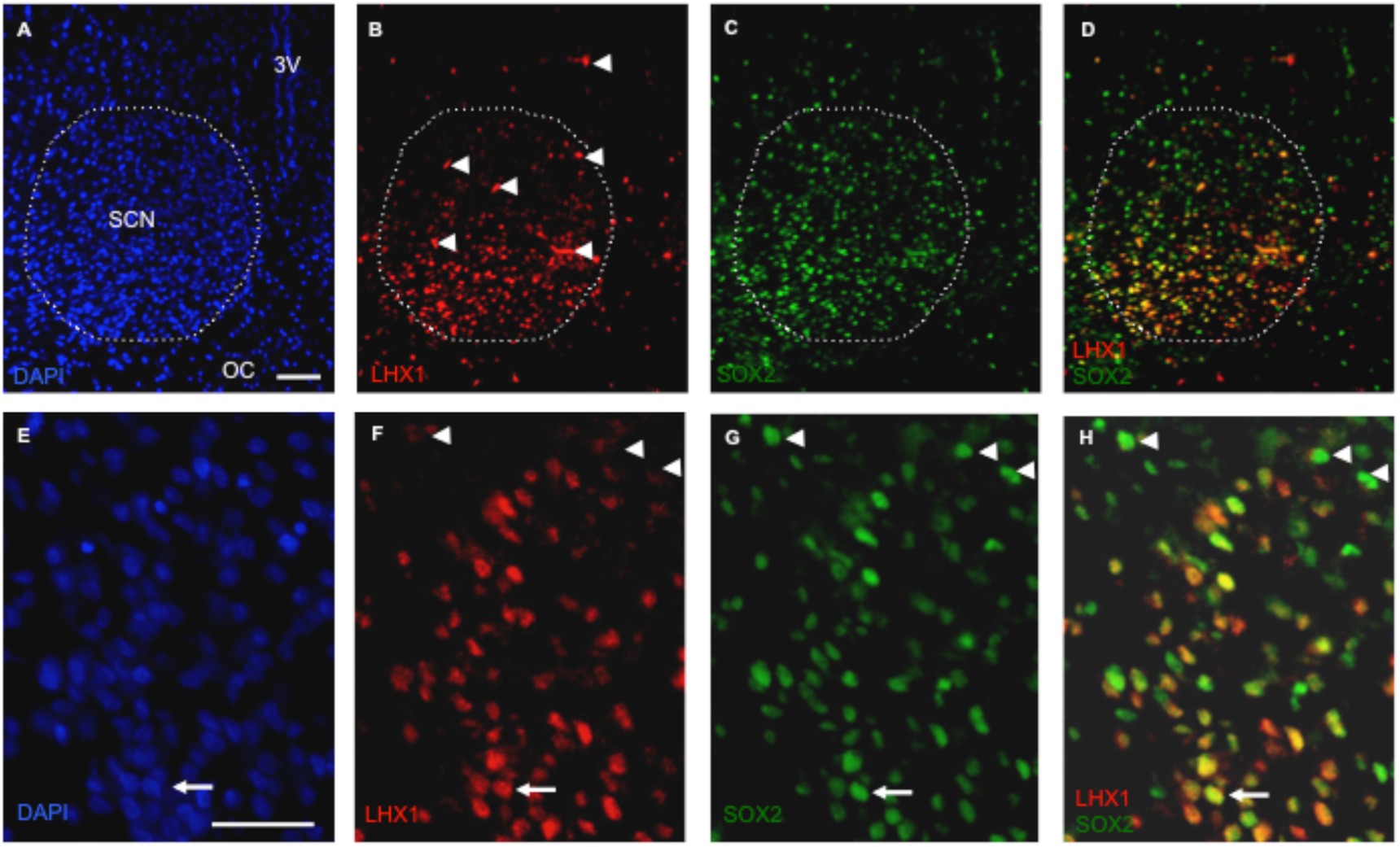
LHX1 is extensively expressed together with SOX2 in the ventromedial region of the rat suprachiasmatic nucleus (SCN), but is largely absent from SOX2+ neurons in the dorsal SCN. Representative fluorescence microscopic images of male PN50 brain showing the distribution of LHX1 and SOX2 immunoreactivity in neurons. **A-D** A restricted distribution of LHX1+ neurons in the ventromedial SCN contrasts with the distribution of SOX2+ neurons that are found across the SCN (dashed line indicates the approximate confines on the SCN). Arrowheads indicate some detection of blood vessels that is observed uniformly across the brain (See Supplemental Figure S10), and does not represent endogenous LHX1. **D-F.** LHX1 is expressed in the nuclear region of SCN neurons, and is extensively co-expressed with SOX2 (one example of co-localization is indicated by the arrow). However, many dorsal SOX2+ neurons (three examples indicated by arrowheads) exhibit minimal/no LHX1 immunoreactivity. Abbreviations: 3V, third ventricle; OC, optic chiasm; DAPI, 4s,6-diamidino-2-phenylindole. Scale bars: 50µm.

## DISCUSSION

Current functional annotation across entire genomes is revealing a complex association between non-coding transcription, and the regulation of coding gene expression (Clark & Blackshaw 2014; Vucicevic et al, 2015; Li et al, 2016; Ntini & Marsico, 2019). Trying to understand this functional interaction is daunting in systems such as the mammalian brain in which there are dynamic changes in gene expression across life (Silberis et al, 2016), but further research is necessary for meaningful understanding of both normal brain function, and the brain dysfunction of neurological and psychiatric diseases. In the current study, an investigation was made into specific, non-coding transcripts across the *Sox2* gene locus with the aim of identifying brain region-specific transcription that may provide insight into the mechanistic basis of adult *Sox2* expression. This analysis of the lncRNA *Sox2ot*, which is transcribed in multiple isoforms from across the *Sox2* locus, and is already implicated in *Sox2* regulation (Messemaker et al, 2018; Shahryari et al, 2015), has revealed novel, region-specific transcripts and patterns of expression. However, the expression profile does not have a distinct, function-oriented, specificity for SCN either in isolation, or when viewed in relation to functionally-annotated *Sox2* enhancers. Rather, subsequent analysis of other regulatory features within the *Sox2* locus, including CTCF sites, indicated that a CTCF-associated non-transcribed enhancer sequence may be, at least partially, permissive for adult brain expression of *Sox2* in the SCN.

Non-coding RNAs are a major component of the transcriptome (Iyer et al, 2015), with temporal and spatial specificity of expression (Reddy et al, 2017; Zhang et al, 2017) that is indicative of cell-specific function. Studies to date indicate roles for lncRNAs in both brain development and disease (Briggs et al, 2015; Parikshak et al, 2016); for *Sox2ot* specifically, a genetic association with anorexia nervosa (Boraska et al, 2014) is one indication of potential roles in nervous system dysfunction. The current analysis of *Sox2ot* expression in adult rat brain has confirmed and extended previous studies (Amaral et al, 2009), confirming both the high level and complex structure of this lncRNA in adult brain (Shahryari et al, 2015; Kadakkuzha et al, 2015; Liu et al, 2016), and, in general, an inverse expression relationship with *Sox2* (see Hoefflin & Carter, 2014). However, despite the current demonstration of low relative levels of one class of *Sox2ot* transcripts in the SOX2-rich OB, the *Sox2:Sox2ot* expression relationship is not uniformly inverse, and consequently the current data is also not consistent with a hypothesized similarity between OB and SCN expression. Rather, the current data indicates diverse, region-specific patterns of *Sox2ot* expression in the adult brain that do not overtly reflect levels of co-expressed *Sox2*. And, secondarily, our findings are consistent with general observations of cell-specific differential splicing (Raj & Blencowe, 2015). Further studies are required to understand the functional relationship between *Sox2ot* and *Sox2* expression in brain that may involve both positive (Xiang et al, 2014) and/or repressive (Messemaker et al, 2018; Spadaro et al, 2015) actions. A recent study (Cheng et al, 2019) has shown that *Sox2ot* expression is up-regulated in the SCN of *Sox2* knockout mice, but only during one phase of the day, indicating that further studies must also address circadian variation in the *Sox2:Sox2ot* relationship. Finally, with respect to some of the (relatively) short variations in *Sox2ot* RNA sequence identified here across brain regions (eg. 4bp deletion in exon7 in cortex), these are representative of recognized splicing variations (tandem alternative splice sites in this case; Szafranski et al, 2014) and should be viewed as potentially functionally important due to the known role of RNA secondary structure in lncRNA function (Clark & Blackshaw 2014).

The current correlative analysis of region-specific *Sox2ot* exon expression and *Sox2* enhancers has shown that many *Sox2ot* exons have a direct correspondence with conserved *Sox2* enhancers (Okamoto et al, 2015), in agreement with general observations (Vucicevic et al, 2015; Li et al, 2016; Hon et al, 2017). As noted, however, no clear relationship was identified between this pattern of lncRNA expression and adult expression of *Sox2* in the SCN. This observation may be partly explained by an incomplete annotation of *Sox2* enhancers – apparent in the current analysis (see Results). Hence, although the extensive studies conducted to date have identified multiple *Sox2* enhancers (27 neuro-sensory), it is apparent that additional enhancers extend beyond the 200kb region analysed by Okamoto and colleagues (2015), consistent with studies of other ‘gene desert’ loci (eg. El-Kasti et al, 2012). For example, the Vista 192 enhancer element that lies on the border of the *Sox2* locus is a likely *Sox2* enhancer due to forebrain-specific activity (enhancer.lbl.gov; see Dickel et al, 2018), and positioning within a genomic context where upstream genes (*Dnajc19* and *Fxr1*) have no brain-enriched expression (GTEx). In confirmation of previous work in other species (Amaral et al, 2009), the current work has shown that exon 1 of rat *Sox2dot* overlaps the isogenic region of this Vista enhancer sequence, and has also shown that the two distal rat exons 2 and 3 are either associated with, or proximal to enhancer chromatin marks. Futhermore, given the multiple additional putative enhancer sequences identified in the current study, and notwithstanding some redundancy (Osterwalder et al, 2018), it is clear that extensive further studies are required to complete the functional annotation of *Sox2* enhancers, and thereafter define their functional association with *Sox2ot* transcripts. Nevertheless, as discussed below, the current study shows that some currently annotated *Sox2* enhancers are both not associated with Sox2ot transcripts, and are also not transcribed at detectable levels.

Further insight into regulatory sequences that direct adult brain expression of *Sox2* was obtained by analysing the relationship between annotated *Sox2* enhancers and CTCF protein associations across the genetic locus. CTCF has multiple roles in addition to the recognized ‘insulator’ activity (Holwerda & de Laat, 2013; Braccioli & de Wit, 2019), and binding to target sites can be either repressive (Lee et al, 2017) or activating (Magbanua et al, 2015) for gene expression. In the current study, a CTCF site was identified that exhibited evidence of specificity for SCN (vs. cortex) in adult brain. This experimental finding can be considered consistent with ENCODE (CTCF ChIP-seq-based) annotation of this site in the mouse genome which reveals relative specificity for embryonic brain (viz. immaturity of gene expression in SCN) compared with adult brain. This adult SCN-active, CTCF site is proximal to a conserved enhancer sequence (denoted U6; Okamoto et al, 2015) that is functional, directing expression to the ventral diencephalon (location of SCN) of chicken, and is highly conserved also in mouse, human and rat (Okamoto et al, 2015). It can therefore be proposed that CTCF acts positively, together with the U6 enhancer sequence, to influence the *Sox2* promoter, and contribute to adult expression of *Sox2* in sub-sets of neurons. This contention is consistent with studies showing that CTCF can stabilize enhancer-promoter interaction (Ren et al, 2017), although the mechanisms involved remain undefined, particularly given recent live-cell imaging studies showing an absence of enhancer proximity during *Sox2* transcription (Alexander et al, 2019). Location of the U6 enhancer within a brain super-enhancer (Fig.6) indicates that U6 is one part of a larger regulatory region that likely has additional, and possibly diverse, functional activity. In this context, annotation of this region in the human genome for selective cingulate gyrus super-enhancer activity (Hnisz et al, 2013), indicates that this genomic region could contribute to a number of distinct regional specificities. Individual cell-type/region-specific regulation could then involve an interaction with other related enhancers, for example the N2 and N3 diencephalic enhancers, in the case of the SCN (Uchikawa et al, 2003; Okamoto et al, 2015).

The current study has not only assembled evidence of a role for U6 enhancer sequence in directing cell-specific *Sox2* expression but also provided evidence of a trans-acting factor that associates with this sequence, namely LHX1, a known regulator of *Sox2* (Inoue et al, 2013; Tam et al, 2016). This evidence is appealing in the light of known roles for LHX1 in the SCN, where it directs aspects of both cell-specific development, and function, in this brain structure (Bedont et al, 2014; 2017). However, subsequent findings here have shown that while LHX1 is certainly abundantly expressed in the rat SCN, it is present in only a (major) subset of SOX2+ve neurons, arguing against a common, and required role for this factor in the maintenance of SCN-wide *Sox2* expression. The biochemical evidence obtained in the current study would be consistent with a developmental involvement of LHX1 in the establishment of *Sox2* expression across the SCN, but absence of a common required role (and expression) in adulthood. Further studies are required to investigate this possibility, and also the different phenotypic attributes of SOX2+ve/LHX1+ve and conversely SOX2/LHX1-ve SCN neurons. Additionally, the likely contribution of other transcription factors to LHX1 function at the U6 enhancer (see Grossman et al, 2017) must also be established.

In summary, the current study has investigated brain region-specific activity at different regulatory features across the *Sox2* locus, and has found novel evidence of region-specific activity that is consistent with a (partial) role in directing adult expression of SOX2 to a specific region of the hypothalamus. This analysis has developed our understanding of the complex logic of gene regulation that is directed through this locus (Okamoto et al, 2015; Sugahara et al, 2018), and posed many questions for further study. The current findings have also highlighted the high, and complex expression pattern of *Sox2ot* in the adult brain, prompting a need for further functional analysis. It is also pertinent to note that the (generally) inverse expression pattern of *Sox2ot* and *Sox2* that arises during the postnatal period in the mammalian brain is one example of the multiple postnatal refinements in neuronal gene expression that are functionally obscure, and could relate to disorders of development. In humans, *Sox2* levels are dynamically regulated during brain development, decreasing acutely during the child (1-4 yrs) to teenager (14-17 yrs) transition, and so forming part of a developmentally regulated set of transcripts that arguably could contribute to neurodevelopmental disorders such as schizophrenia (Jaffe et al, 2015). The latter dataset (Jaffe et al, 2015) also shows that particular *Sox2ot* exons exhibit a converse, increase in expression across postnatal development into adulthood, indicating again a possible relatedness of these events (the postnatal increase in *Sox2ot* expression may be causally linked to the fall in *Sox2* expression). Consequently, a greater understanding of the relationship between *Sox2* and *Sox2ot* expression could also be valuable for the elucidation of molecular phenotypes that relate to disease.

## CONFLICT OF INTERESTS

The author declares that the research was conducted in the absence of any commercial or financial relationships that could be construed as a potential conflict of interest.

## FUNDING

This study was supported by the Cardiff School of Biosciences, Cardiff University, UK.

## ACKNOWLEDGEMENTS

Dr Jeff Davies (Swansea University, UK) is thanked for the HT-22 cells. A number of Cardiff University undergraduate project students (Christopher Grubb, Stuart Yule, Isobel Sutherland) contributed to preliminary studies on the Sox2ot gene, and their contribution is acknowledged.

