## Supplemental data for "Integrated analysis of different non-coding features across the *Sox2* locus implicates a diencephalic enhancer in adult brain expression"

**S1.**

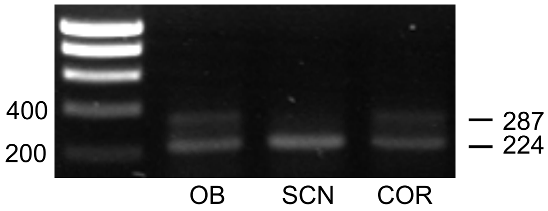


Fig. S1. Differential expression of *Grin1* exon 4 in suprachiasmatic nucleus (SCN), as compared with olfactory bulb (OB) and brain cortex (COR). Representative agarose gel electrophoresis image of RT-PCR analysis using primers directed part of *Grin1* mRNA. Note the virtual exclusion of the exon 4-containing (287bp) product in SCN alone, but similar expression of the exon 4-excluding (224bp) product in all three PCR reactions. Numbers indicate amplicon sizes, and size ladder bands (left) in bp.

**S2. Rat brain Sox2ot & Sox2dot cDNA sequences, and exon structure.**
Dinucleotides that immediately flank exons were taken from genomic database sequence (Rnor 6.0), and are shown in lower case, red font.

#### F8R3 (722bp, cortex)

CCTGCACAGGGCTGGCTGTGGCAGGGGACACTCTCCTGCTGGCTCTCTCTGACCACACATTAGCTGGCT
CCCATCCTCAGGTGCACGGCCACACCCTTGCACCCAACCTTGATCCTCTGATGGGGAAGGTCCCAGGAC
TTGCCAGCTGGGACAGGCCTCACCCCCAAGCGCTCTATGCAGTACTGAGAAGCAAACCTGACAGATTTC
TCCAGAAAAGTGGCCATCCATGGAATGAGTGAAATACTCTTTCTATTCCAGGGATTGCAGAGGCAAAGC
TAGGCTAGGTCTTGGAGGCTGGTGGAAGGCGGTGCGGGTAAGGCAGGACGCTGATGGGAGAGACTGGTC
GAAGAAAAGCCGAAATGGATTCTCCGTGCCTTGGATGGAGGAAGAGGGGGGAAGTGCGAGCTCCTTCAA
CTCGTTCTGTCCGGTGAGGAGTGATCCAGCTTGGGCTGACAAGAGGCTCCGGAGCCTTCATGTTCAGTG
CTTTTTTACCTGCCAATCAAACTGCTACAAGACAACACCCTGCTCTGGCATGGACAACCTAGAAATTAT
ATTCTCTTCAAGATCCATGAAAGAGAACCTGACTGCTTGCTTTGGTCTGAGAGTTCAGACATCTCTTCT
CTTAAGAAAGTCTTCATTTCCGTTTGCTGAGTCCTCCTGTTTTGTCATCGCTGGGCCCTGTAAGAAGTT
TGCCAGGAGGTCCCCAGAAGTAGAAGGACCAT

#### Exon structure

tcCCTGCACAGGGCTGGCTGTGGCAGGGGACACTCTCCTGCTGGCTCTCTCTGACCACACATTAGCTGG
CTCCCATCCTCAGGTGCACGGCCACACCCTTGCACCCAACCTTGATCCTCTGATGGGGAAGGTCCCAGG
ACTTGCCAGCTGGGACAGGCCTCACCCCCAAGCGCTCTATGCAGTACTGAGAAGgt

agCAAACCTGACAGATTTCTCCAGAAAAGTGGCCATCCATGGAATGAGTGAAATACTCTTTCTATTCCA
gt

agGGGATTGCAGAGGCAAAGCTAGGCTAGGTCTTGGAGGCTGGTGGAAGGCGGTGCGGGTAAGGCAGGA
CGCTGATGGGAGAGACTGGTCGAAGAAAAGCCGAAATGGATTCTCCGTGCCTTGGATGGAGGAAGAGGG
GGGAAGTGCGAGCTCCTTCAACTCGTTCTGTCCGGTGAGGAGTGATCCAGCTTGGGCTGACAAGAGGCT
CCGGAGCCTTCATGTTCAGTGCTTTTTTACCTGCCAATCAAACTGCTACAAGACAACACCCTGCTCTGG
CATGGACgt

agAACCTAGAAATTATATTCTCTTCAAGATCCATGAAAGAGAACCTGACTGCTTGCTTTGGTCTGAGAG
TTCAGACATCTCTTCTCTTAAGAAAGTCTTCATTTCCGTTTGCTGAGTCCTCCTGTTTTGTCATCGCTG
GGCCCTGTAAGAAGTTTGCCAGGAGGTCCCCAGAAGTAGAAGGACCAT

#### F8R3 (726bp, SCN and olfactory bulb)

CCTGCACAGGGCTGGCTGTGGCAGGGGACACTCTCCTGCTGGCTCTCTCTGACCACACATTAGCTGGCT
CCCATCCTCAGGTGCACGGCCACACCCTTGCACCCAACCTTGATCCTCTGATGGGGAAGGTCCCAGGAC
TTGCCAGCTGGGACAGGCCTCACCCCCAAGCGCTCTATGCAGTACTGAGAAGCAAACCTGACAGATTTC
TCCAGAAAAGTGGCCATCCATGGAATGAGTGAAATACTCTTTCTATTCCAGGGATTGCAGAGGCAAAGC
TAGGCTAGGTCTTGGAGGCTGGTGGAAGGCGGTGCGGGTAAGGCAGGACGCTGATGGGAGAGACTGGTC
GAAGAAAAGCCGAAATGGATTCTCCGTGCCTTGGATGGAGGAAGAGGGGGGAAGTGCGAGCTCCTTCAA
CTCGTTCTGTCCGGTGAGGAGTGATCCAGCTTGGGCTGACAAGAGGCTCCGGAGCCTTCATGTTCAGTG
CTTTTTTACCTGCCAATCAAACTGCTACAAGACAACACCCTGCTCTGGCATGGACATAGAACCTAGAAA
TTATATTCTCTTCAAGATCCATGAAAGAGAACCTGACTGCTTGCTTTGGTCTGAGAGTTCAGACATCTC
TTCTCTTAAGAAAGTCTTCATTTCCGTTTGCTGAGTCCTCCTGTTTTGTCATCGCTGGGCCCTGTAAGA
AGTTTGCCAGGAGGTCCCCAGAAGTAGAAGGACCAT

#### Exon structure

tcCCTGCACAGGGCTGGCTGTGGCAGGGGACACTCTCCTGCTGGCTCTCTCTGACCACACATTAGCTGG
CTCCCATCCTCAGGTGCACGGCCACACCCTTGCACCCAACCTTGATCCTCTGATGGGGAAGGTCCCAGG
ACTTGCCAGCTGGGACAGGCCTCACCCCCAAGCGCTCTATGCAGTACTGAGAAGgt

agCAAACCTGACAGATTTCTCCAGAAAAGTGGCCATCCATGGAATGAGTGAAATACTCTTTCTATTCCA
gt

agGGGATTGCAGAGGCAAAGCTAGGCTAGGTCTTGGAGGCTGGTGGAAGGCGGTGCGGGTAAGGCAGGA
CGCTGATGGGAGAGACTGGTCGAAGAAAAGCCGAAATGGATTCTCCGTGCCTTGGATGGAGGAAGAGGG
GGGAAGTGCGAGCTCCTTCAACTCGTTCTGTCCGGTGAGGAGTGATCCAGCTTGGGCTGACAAGAGGCT
CCGGAGCCTTCATGTTCAGTGCTTTTTTACCTGCCAATCAAACTGCTACAAGACAACACCCTGCTCTGG
CATGGACgt

agATAGAACCTAGAAATTATATTCTCTTCAAGATCCATGAAAGAGAACCTGACTGCTTGCTTTGGTCTG
AGAGTTCAGACATCTCTTCTCTTAAGAAAGTCTTCATTTCCGTTTGCTGAGTCCTCCTGTTTTGTCATC
GCTGGGCCCTGTAAGAAGTTTGCCAGGAGGTCCCCAGAAGTAGAAGGACCAT

## F1BR3 (743bp)

ACAGAGGAGAAGCGGCTGAGGAGAGGAAGGAAGGAGTCTATCACATCATCTCTCTGTCCACAGGCTTTC
CTTGGGGCTTATGAAGATTTCTTCGCACCACCCCTCTTCACCCCTTGGAACCGGAAGGATGTGGATAGA
TGTCTTAAGAACAAAATGAACTGGAATTTTTTTGAGAGGAAGAACAGCATGAGACACTTACTTGTGTCC
TCGGAAGCCTGCACAGATATTCCCATCCCCTCTTCCCGAGTCACCACCTACTTCAGTGGAATATAAGGG
GATTGCAGAGGCAAAGCTAGGCTAGGTCTTGGAGGCTGGTGGAAGGCGGTGCGGGTAAGGCAGGACGCT
GATGGGAGAGACTGGTCGAAGAAAAGCCGAAATGGATTCTCCGTGCCTTGGATGGAGGAAGAGGGGGGA
AGTGCGAGCTCCTTCAACTCGTTCTGTCCGGTGAGGAGTGATCCAGCTTGGGCTGACAAGAGGCTCCGG
AGCCTTCATGTTCAGTGCTTTTTTACCTGCCAATCAAACTGCTACAAGACAACACCCTGCTCTGGCATG
GACATAGAACCTAGAAATTATATTCTCTTCAAGATCCATGAAAGAGAACCTGACTGCTTGCTTTGGTCT
GAGAGTTCAGACATCTCTTCTCTTAAGAAAGTCTTCATTTCCGTTTGCTGAGTCCTCCTGTTTTGTCAT
CGCTGGGCCCTGTAAGAAGTTTGCCAGGAGGTCCCCAGAAGTAGAAGGACCAT

### Exon structure

gcACAGAGGAGAAGCGGCTGAGGAGAGGAAGGAAGGAGTCTATCACATCATCTCTCTGTCCACAGGCTT
TCCTTGGGGCTTATGAAGgt

agATTTCTTCGCACCACCCCTCTTCACCCCTTGGAACCGGAAGGATGTGGATAGATGTCTTAAGAACAg
t
agAAATGAACTGGAATTTTTTTGAGAGGAAGAACAGCATGAGACACTTACTTGTGTCCTCGGAAGCCTG
CACAGATATTCCCATCCCCTCTTCCCGAGTCACCACCTACTTCAGTGGAATATAAGgt

agGGGATTGCAGAGGCAAAGCTAGGCTAGGTCTTGGAGGCTGGTGGAAGGCGGTGCGGGTAAGGCAGGA
CGCTGATGGGAGAGACTGGTCGAAGAAAAGCCGAAATGGATTCTCCGTGCCTTGGATGGAGGAAGAGGG
GGGAAGTGCGAGCTCCTTCAACTCGTTCTGTCCGGTGAGGAGTGATCCAGCTTGGGCTGACAAGAGGCT
CCGGAGCCTTCATGTTCAGTGCTTTTTTACCTGCCAATCAAACTGCTACAAGACAACACCCTGCTCTGG
CATGGACgt

agATAGAACCTAGAAATTATATTCTCTTCAAGATCCATGAAAGAGAACCTGACTGCTTGCTTTGGTCTG
AGAGTTCAGACATCTCTTCTCTTAAGAAAGTCTTCATTTCCGTTTGCTGAGTCCTCCTGTTTTGTCATC
GCTGGGCCCTGTAAGAAGTTTGCCAGGAGGTCCCCAGAAGTAGAAGGACCAT

## F1BR3 (512bp)

ACAGAGGAGAAGCGGCTGAGGAGAGGAAGGAAGGAGTCTATCACATCATCTCTCTGTCCACAGGCTTTC
CTTGGGGCTTATGAAGATTTCTTCGCACCACCCCTCTTCACCCCTTGGAACCGGAAGGATGTGGATAGA
TGTCTTAAGAACAAAATGAACTGGAATTTTTTTTGAGAGGGGGGAAGTGCGAGCTCCTTCAACTCGTTC
TGTCCGGTGAGGAGTGATCCAGCTTGGGCTGACAAGAGGCTCCGGAGCCTTCATGTTCAGTGCTTTTTT
ACCTGCCAATCAAACTGCTACAAGACAACACCCTGCTCTGGCATGGACATAGAACCTAGAAATTATATT
CTCTTCAAGATCCATGAAAGAGAACCTGACTGCTTGCTTTGGTCTGAGAGTTCAGACATCTCTTCTCTT
AAGAAAGTCTTCATTTCCGTTTGCTGAGTCCTCCTGTTTTGTCATCGCTGGGCCCTGTAAGAAGTTTGC
CAGGAGGTCCCCAGAAGTAGAAGGACCAT

### Exon structure

gcACAGAGGAGAAGCGGCTGAGGAGAGGAAGGAAGGAGTCTATCACATCATCTCTCTGTCCACAGGCTT
TCCTTGGGGCTTATGAAGgt

agATTTCTTCGCACCGCCCCTCTTCACCCCTTGGAACCGGAAGGATGTGGATAGATGTCTTAAGAACAg
t

agAAATGAACTGGAATTTTTTTTGAGAGga

agGGGGGAAGTGCGAGCTCCTTCAACTCGTTCTGTCCGGTGAGGAGTGATCCAGCTTGGGCTGACAAGA
GGCTCCGGAGCCTTCATGTTCAGTGCTTTTTTACCTGCCAATCAAACTGCTACAAGACAACACCCTGCT
CTGGCATGGACgt

agATAGAACCTAGAAATTATATTCTCTTCAAGATCCATGAAAGAGAACCTGACTGCTTGCTTTGGTCTG
AGAGTTCAGACATCTCTTCTCTTAAGAAAGTCTTCATTTCCGTTTGCTGAGTCCTCCTGTTTTGTCATC
GCTGGGCCCTGTAAGAAGTTTGCCAGGAGGTCCCCAGAAGTAGAAGGACCAT

**F1BR3 (419bp)**
ACAGAGGAGAAGCGGCTGAGGAGAGGAAGGAAGGATGTGGATAGATGTCTTAAGAACAAAATGAACTGG
AATTTTTTTTGAGAGGGGGGAAGTGCGAGCTCCTTCAACTCGTTCTGTCCGGTGAGGAGTGATCCAGCT
TGGGCTGACAAGAGGCTCCGGAGCCTTCATGTTCAGTGCTTTTTTACCTGCCAATCAAACTGCTACAAG
ACAACACCCTGCTCTGGCATGGACATAGAACCTAGAAATTATATTCTCTTCAAGATCCATGAAAGAGAA
CCTGACTGCTTGCTTTGGTCTGAGAGTTCAGACATCTCTTCTCTTAAGAAAGTCTTCATTTCCGTTTGC
TGAGTCCTCCTGTTTTGTCATCGCTGGGCCCTGTAAGAAGTTTGCCAGGAGGTCCCCAGAAGTAGAAGG
ACCAT

**Exon structure**
gcACAGAGGAGAAGCGGCTGAGGAGAGGAAGGAAGga

agGATGTGGATAGATGTCTTAAGAACAgt

agAAATGAACTGGAATTTTTTTTGAGAGga

agGGGGGAAGTGCGAGCTCCTTCAACTCGTTCTGTCCGGTGAGGAGTGATCCAGCTTGGGCTGACAAGA
GGCTCCGGAGCCTTCATGTTCAGTGCTTTTTTACCTGCCAATCAAACTGCTACAAGACAACACCCTGCT
CTGGCATGGACgt

agATAGAACCTAGAAATTATATTCTCTTCAAGATCCATGAAAGAGAACCTGACTGCTTGCTTTGGTCTG
AGAGTTCAGACATCTCTTCTCTTAAGAAAGTCTTCATTTCCGTTTGCTGAGTCCTCCTGTTTTGTCATC
GCTGGGCCCTGTAAGAAGTTTGCCAGGAGGTCCCCAGAAGTAGAAGGACCAT

**S3. Peptide sequence encoded by ORFs in rat brain Sox2ot/Sox2dot RNAs (722/726 & 743 bp amplicons).** 114, and 91 amino acid rat sequences are shown, and, for comparison, a homologous, database-annotated, 106 amino acid mouse sequence

Rat (722bp):
MEEEGGSASSFNSFCPVRSDPAWADKRLRSLHVQCFFTCQSNCYKTTPCSGMD
NLEIIFSSRSMKENLTACFGLRVQTSLLLRKSSFPFAESSCFVIAGPCKKFARRSPEVEGP

Rat (726 & 743bp):
MEEEGGSASSFNSFCPVRSDPAWADKRLRSLHVQCFFTCQSNCYKTTPCSGMD
IEPRNYILFKIHEREPDCLLWSESSDISSLKKVFISVC

Mouse: >uc012cor.1 (Sox2ot) length=106
MEKEGGSASSFNSFCPVRSDPAWADERLRSLHVQCFLPANQTATRQHPDLAWSPRVQACS
PKSENLCSWYTQKRSQATDILQKKALNSEIEPRNCTFLKIRLRESG

**S4. Analysis of consensus ribonucleoprotein binding sites in Sox2dot sequence.** Highlighted sequences are consensus sites: blue, NOVA (YCAY); green, RBFOX ([U]GCAUG). Sequences in red text are regions deleted in some Sox2dot isoforms.

**F8R3 (726bp OB/SCN)**

CCTGCACAGGGCTGGCTGTGGCAGGGGACACTCTCCTGCTGGCTCTCTCTGACCACACATTAGCTGGCT
CCCATCCTCAGGTGCACGGCCACACCCTTGCACCCAACCTTGATCCTCTGATGGGGAAGGTCCCAGGAC
TTGCCAGCTGGGACAGGCCTCACCCCCAAGCGCTCTATGCAGTACTGAGAAGCAAACCTGACAGATTTC
TCCAGAAAAGTGGCCATCCATGGAATGAGTGAAATACTCTTTCTATTCCAGGGATTGCAGAGGCAAAGC
TAGGCTAGGTCTTGGAGGCTGGTGGAAGGCGGTGCGGGTAAGGCAGGACGCTGATGGGAGAGACTGGTC
GAAGAAAAGCCGAAATGGATTCTCCGTGCCTTGGATGGAGGAAGAGGGGGGAAGTGCGAGCTCCTTCAA
CTCGTTCTGTCCGGTGAGGAGTGATCCAGCTTGGGCTGACAAGAGGCTCCGGAGCCTTCATGTTCAGTG
CTTTTTTACCTGCCAATCAAACTGCTACAAGACAACACCCTGCTCTGGCATGGACATAGAACCTAGAAA
TTATATTCTCTTCAAGATCCATGAAAGAGAACCTGACTGCTTGCTTTGGTCTGAGAGTTCAGACATCTC
TTCTCTTAAGAAAGTCTTCATTTCCGTTTGCTGAGTCCTCCTGTTTTGTCATCGCTGGGCCCTGTAAGA
AGTTTGCCAGGAGGTCCCCAGAAGTAGAAGGACCAT

### F1BR3 (743, SCN)

ACAGAGGAGAAGCGGCTGAGGAGAGGAAGGAAGGAGTCTATCACATCATCTCTCTGTCCACAGGCTTTC
CTTGGGGCTTATGAAGATTTCTTCGCACCACCCCTCTTCACCCCTTGGAACCGGAAGGATGTGGATAGA
TGTCTTAAGAACAAAATGAACTGGAATTTTTTTGAGAGGAAGAACAGCATGAGACACTTACTTGTGTCC
TCGGAAGCCTGCACAGATATTCCCATCCCCTCTTCCCGAGTCACCACCTACTTCAGTGGAATATAAGGG
GATTGCAGAGGCAAAGCTAGGCTAGGTCTTGGAGGCTGGTGGAAGGCGGTGCGGGTAAGGCAGGACGCT
GATGGGAGAGACTGGTCGAAGAAAAGCCGAAATGGATTCTCCGTGCCTTGGATGGAGGAAGAGGGGGGA
AGTGCGAGCTCCTTCAACTCGTTCTGTCCGGTGAGGAGTGATCCAGCTTGGGCTGACAAGAGGCTCCGG
AGCCTTCATGTTCAGTGCTTTTTTACCTGCCAATCAAACTGCTACAAGACAACACCCTGCTCTGGCATG
GACATAGAACCTAGAAATTATATTCTCTTCAAGATCCATGAAAGAGAACCTGACTGCTTGCTTTGGTCT
GAGAGTTCAGACATCTCTTCTCTTAAGAAAGTCTTCATTTCCGTTTGCTGAGTCCTCCTGTTTTGTCAT
CGCTGGGCCCTGTAAGAAGTTTGCCAGGAGGTCCCCAGAAGTAGAAGGACCAT

### S5. Conservation of U6 enhancer sequence in rat.

#### Human

>hg19_dna range=chr3:181343682-181344425
AGAGGTGAAATATTGCTTTTACTGTTTTCAGATAATAGTGACTGTTGTTG
CAAATTATAAAATTTAATTAGCTATATAAATGCCAGAGGAACAAATAATG
TGCTATTGTCTTGATATCTTTCCAGAAATTAGTCTTTTAAAATAAATTTG
CCATATTAATCATCTAAAAATATTTTTGATCACTGAATTAATAGTTTGCA
GCCTGTTTTCAGGTACTCATTGCCATAGGTGATTGCTTCTTAATTACTAC
ATGGGCTTTTTCTATTCACTCATATTAATTCATGTTTTCTGGCACAGTCC
CAAAAGGCAGCTTGCCCTATATCTTCTAATTACTTCTATAATTGAATTTG
CCCACTTTTTCTAACCTATTTTTTAAACTGACTGTGTTTGTTCATGGTTT
ATTGTGTCTTTTTTGTGTCAGTTCTCTGCAAGCAGAATGTACATTGATTC
TTTCCTCGTTTCTCATTAATTTTCAAGAGCTTTGCTGCATGTCTGTTTTT
TCAGTCTGTCAGACCTATTGCTAGACTCCCCCCCTCAAAAAAAGGCTCTT
ATTATTAAAGGTTAGGTAAAATAGACTGAAGTTAATTAATTCATTAGTAA
CTCTGAGCCAATTACTGAAGCCTTCTCTTATTTATATACTTGTTAACAAG
ACAGAAATAAGAAAGGCCAATTACAGGCTCATGTCCTTCAGAAGTTCAGC
AAACCC

#### Rat

>rn5_dna range=chr2:140725908-140726750
AGAGGTGAAATACTGCCTTCACTGTTTTCAGATAATAATGACTGTTGTTGCA
AATTATAAAATTTAATTAGCTATATAAATGCCAGAAGAACAAACGGTATG
CCATTGTCTTGATATCTTTCCAGAAATTAGTCTTTTAAAAATAAATTTGC
CATATTAATCATCTAAAAATATTTTTGATCACTGAATTAATGGTTTTAGG
CCTGTTTTCAGGTGCTCATTGCTATAGGTGATTGCTTCTTAATTACTACA
TGGGCTCTCTTTTCACTCATATTAATTCATGTTTGCTGGCACAGTCCCAA
AGGCAGCCTGCCCTGGATCTTCTAATTACTTCTATAATTGAATTTGCCCA
CTTTTTCTAACCTATTTTTTAAACTGACTGTGTTTGTCCCAAGCTTATTG
TGTCTTTTTTGTGTCAGTTCCCCGCAAGCAGAGAGTACATCGATCCTTTC
CTCGTTTCTCATTAATTTTCAAGAGCTTGGCCGCACGTCTGGTTTTTTTT
TTTTTTTTTTTCTTCAGTCCGTCAGACCTATTGCAAAATGGAAGAAAAAA
GAAGGAAAGAAAGGACGGAAAAAAAGAGAAAAAAAAGGGACTTTTATTATT

```
AAAGGTTAGGTAAAATAGACTGAAGTTAATTAATTCATTAGTAACCCTGA
GCCAATTACTGAAGCCTTCTCTTATTTATATACTTGGTAACAAGACAGAA
ATGAGAAAGGCCAATTACAGGCACATGTCCTTCACAAGTTCAGCCACCTC
```

**Mouse**

```
>mm9_dna range=chr3:34476488-34477365
AGAGGTGAAATATTGCCTTCACTGTTTTCAGATAATAATG
ACTGTTGTTGCAAATTATAAAATTTAATTAGCTATATAAATTCCAGAAGA
ACAAACGGTATGCCATTGTCTTGATATCTTTCCAGAAATTAGTCTTTTAA
AAAATAAATTTGCCATATTAATCATCTAAAAATATTTTTGATCACTGAAT
TAATGGTTTTAGGCCTGTTTTCAGGTGCTCATTGCTATAGGTGATTGCTT
CTTAATTACTACATGGACTCTCTATTCACTCATATTAATTCATGTTTGCT
GGCACAGTCCCAGAGGCAGCCTGCCCTGGATCTTCTAATTACTTCTATAA
TTGAATTTGCCCACTTTTTCTAACCTATTTTTTAAACTGACTGTGTTTGT
CCCAAGCTTATTGTGTCTTTTTTGTGTCAGTTCCCTGCAAGCAGAGTGTA
CATCGATCCTTTCCTCGTTTCTCATTAATTTTCAAGAGCTTGGCCGCATG
TCGGTTTTTTTTTTTTTTTCAGTCCGTCAGACCTATTGCGAAACGGAAAAG
AAAAAAAAAAAAAAAGAAAGAAAGAAAGAAAGGGAAAGAGGAAAAAGGGAA
AAAAGGACTTTTATTATTAAAGGTTAGGTAAAATAGACTGAAGTTAATTA
ATTCATTAGTAACCCTGAGCCAATTACTGAAGCCTTCTCTTATTTATATA
CTTGTTAACAAGACAGAAACGAGAAAGGCCAATTACAGGCACATGCCCTT
CACAACTTCAGCCCTCTC
```

**CLUSTAL O(1.2.1) multiple sequence alignment of U6 enhancer**

Highlighted sequences (eg. ATTTAATTAG) are Lhx3 consensus sites. Note these 3 sites also conform to the Lhx1 consensus core site: T/c/a/g-T/c/a-A/c/g-A/t-T/a-T/c/g-A/g/t-A/g/t/c as defined for mouse in Jaspar2016 (Mathelier et al, 2016).

```
human      AGAGGTGAAATATTGCTTTTACTGTTTTCAGATAATAGTGACTGTTGTTGCAAATTATAA
rat        AGAGGTGAAATACTGCCTTCACTGTTTTCAGATAATAATGACTGTTGTTGCAAATTATAA
mouse      AGAGGTGAAATATTGCCTTCACTGTTTTCAGATAATAATGACTGTTGTTGCAAATTATAA
           ************ *** ** ***************** **********************

human      AATTTAATTAGCTATATAAATGCCAGAGGAACAAATAATGTGCTATTGTCTTGATATCTT
rat        AATTTAATTAGCTATATAAATGCCAGAAGAACAAACGGTATGCCATTGTCTTGATATCTT
mouse      AATTTAATTAGCTATATAAATTCCAGAAGAACAAACGGTATGCCATTGTCTTGATATCTT
           ********************* ***** *******   * *** ****************

human      TCCAGAAATTAGTCTTTTA--AAATAAATTTGCCATATTAATCATCTAAAAATATTTTTG
rat        TCCAGAAATTAGTCTTTTAA-AAATAAATTTGCCATATTAATCATCTAAAAATATTTTTG
mouse      TCCAGAAATTAGTCTTTTAAAAAATAAATTTGCCATATTAATCATCTAAAAATATTTTTG
           *******************  ***************************************

human      ATCACTGAATTAATAGTTTGCAGCCTGTTTTCAGGTACTCATTGCCATAGGTGATTGCTT
rat        ATCACTGAATTAATGGTTTTAGGCCTGTTTTCAGGTGCTCATTGCTATAGGTGATTGCTT
mouse      ATCACTGAATTAATGGTTTTAGGCCTGTTTTCAGGTGCTCATTGCTATAGGTGATTGCTT
           ************** ****   ************** ******** **************

human      CTTAATTACTACATGGGCTTTTTCTATTCACTCATATTAATTCATGTTTTCTGGCACAGT
rat        CTTAATTACTACATGGGCTCTC--TTTTCACTCATATTAATTCATGTTTGCTGGCACAGT
mouse      CTTAATTACTACATGGACTCTC--TATTCACTCATATTAATTCATGTTTGCTGGCACAGT
           **************** ** *   * *********************** **********

human      CCCAAAAGGCAGCTTGCCCTATATCTTCTAATTACTTCTATAATTGAATTTGCCCACTTT
rat        CCC-AAAGGCAGCCTGCCCTGGATCTTCTAATTACTTCTATAATTGAATTTGCCCACTTT
mouse      CCC-AGAGGCAGCCTGCCCTGGATCTTCTAATTACTTCTATAATTGAATTTGCCCACTTT
           *** * ******* ******  **************************************

human      TTCTAACCTATTTTTTAAACTGACTGTGTTTGTTCATGGTTTATTGTGTCTTTTTTGTGT
rat        TTCTAACCTATTTTTTAAACTGACTGTGTTTGTCCCAAGCTTATTGTGTCTTTTTTGTGT
mouse      TTCTAACCTATTTTTTAAACTGACTGTGTTTGTCCCAAGCTTATTGTGTCTTTTTTGTGT
           ********************************* *   * ********************
```

```
human      CAGTTCTCTGCAAGCAGAATGTACATTGATTCTTTCCTCGTTTCTCATTAATTTTCAAGA
rat        CAGTTCCCCGCAAGCAGAGAGTACATCGATCCTTTCCTCGTTTCTCATTAATTTTCAAGA
mouse      CAGTTCCCTGCAAGCAGAGTGTACATCGATCCTTTCCTCGTTTCTCATTAATTTTCAAGA
           ****** * *********  ****** *** *****************************

human      GCTTTGCTGCATGTCTGTTTTTTCAGTCTGTC-----------------AGACCTATTGC
rat        GCTTGGCCGCACGTCTGGTTTTTTTTTTTTTTTTTCTTCAGTCCGTCAGACCTATTGC
mouse      GCTTGGCCGCATGTCGGTTTTTTTTTTTTTTTCAGT---------CCGTCAGACCTATTGC
           **** ** *** *** * *****    * * *                 ***********

human      TAGAC----------------------------------------------TCCCCCCCTC
rat        AAAATGGAA--------------GAAAAAAGAAGGAAAGAAAGGACGGAAAAAAAGAGAA
mouse      GAAACGGAAAAGAAAAAAAAAAAAAAAGAAAGAAAGAAAGAAAGGGAAAGAGGAAAAAGGG
            * *

human      AAAAAAAGGCTCTTATTATTAAAGGTTAGGTAAAATAGACTGAAGTTAATTAATTCATTA
rat        AAAAAGGGACTTTTATTATTAAAGGTTAGGTAAAATAGACTGAAGTTAATTAATTCATTA
mouse      AAAAAAGGACTTTTATTATTAAAGGTTAGGTAAAATAGACTGAAGTTAATTAATTCATTA
           *****  * ** ************************************************

human      GTAACTCTGAGCCAATTACTGAAGCCTTCTCTTATTTATATACTTGTTAACAAGACAGAA
rat        GTAACCCTGAGCCAATTACTGAAGCCTTCTCTTATTTATATACTTGGTAACAAGACAGAA
mouse      GTAACCCTGAGCCAATTACTGAAGCCTTCTCTTATTTATATACTTGTTAACAAGACAGAA
           ***** **************************************** *************

human      ATAAGAAAGGCCAATTACAGGCTCATGTCCTTCAGAAGTTCAGCAAACCC
rat        ATGAGAAAGGCCAATTACAGGCACATGTCCTTCACAAGTTCAGCC-----
mouse      ACGAGAAAGGCCAATTACAGGCACATGCCCTTCACAACTTCAGCCCTCTC
           *  ******************* **** ****** ** ***
```

## S6.

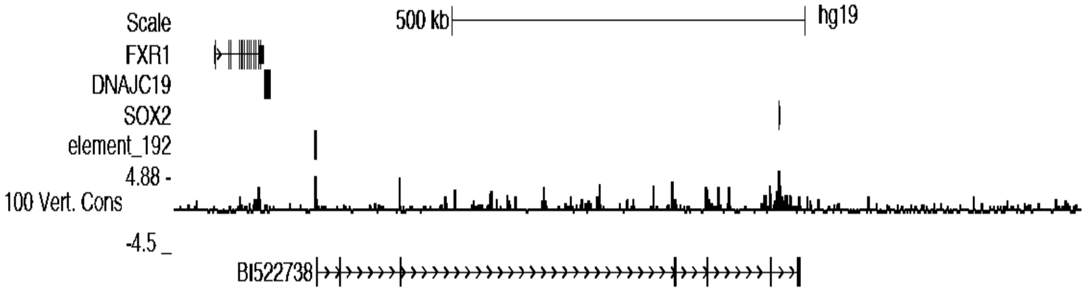


Fig. S6. UCSC genome browser image (Human, hg19) showing that rat Sox2dot exon 1 (conserved in exon 1 of the human EST BI522738) is located within a Vista enhancer sequence (element192). 100 Vert. Cons = Placental mammal basewise conservation (phyloP).

**S7. Additional EST sequences mapping to the Sox2 enhancer region.**

Mouse mm10:

Gencode lincRNA, Gm38505, ENSMUST00000174005.2, antisense, which also maps to ESTs BU936801 and eg. CV308096, maps to mouse Chr 3: 34298172-34351712

Gencode lincRNA, Gm43208-201 ENSMUST00000198924.1, sense, also maps to EST CR515803, maps to Chr 3: 34416495-34418162

Gencode lincRNA, Gm20515-201 ENSMUST00000174440.1, antisense, no ESTs, maps to Chr 3: 34427373-34435097

Gencode lincRNA, Gm29135-201 ENSMUST00000188411.1, antisense, no EST, maps to Chr 3: 34481428-34482297. Partially overlaps U17.

Gencode lincRNA, Gm42695-201 ENSMUST00000199445.1, antisense, no EST, maps to 34565265-34572202

Gencode lincRNA, Gm43207-201 ENSMUST00000199244.1, antisense, EST AI595244, maps to Chromosome 3: 34,598,519-34,602,248. Partially overlaps U4.

**S8. Identification of consensus CTCF sites within CTCF ChIP-Seq regions.**
Sequences were searched using Matinspector (Genomatix)

**1. U6-associated site (labeled 4, Fig. 4A)**

Mouse: sequence extracted from genome browser

```
>mm9_dna range=chr3:34474962-34475770
ATAAAAGGTTATTGTCTTTTCTGTTTTGCTTCGTTTTTTTACTTTGGTTG
TATTGCTGCACGTTATTTTATGGCATGGTCTCATATAGTTCAGGCTTTCT
TCAAATTCCCTTTCTTGTAGGCACGGCTGGCCTTGAACTTTCCATACTCC
CTCCCGCCTAACCTCCAGAGTGCTCAGCTTACAGGTGTCTGCCACGTACT
CTAACCTTTGTCTGCTTTATTCTCTGCATCGCCTCCATCGTTCCCTTCCT
CTCTATTAGGGTGCCTGCCTTCTCTACCCCCTTGTAAAGAGCCTGGGTGT
TCTTTCCAACCAGTGGAGGCGCTGTTGGCAGCCCCATGCTTGCTCTCTCG
CGGTTGACTTTACATGGTGAGTGGGCCCTGTTCAGACCTCTCTAAGCTCT
GCTCTCTTCACTAGTCGTGCCGCCTCTGGGCAGCCCCTGCGCCTGCTCTA
CCTGGGCCTGGCTTCTTAGCCTCAAGCTCACTCAGGGAGTATGAGCCCTG
GTGCATACACACTTCTTGCCCATTTTCTCTCCCAGAATACCCAGTTTGTC
TTTAGAATTAAAAGCTAGAATGGCAACAATGAAATCAGCTTCAGAAAGTA
GCTTGAATGCCTTTCATTTTCTGTTTTTGTCTATGCCAGTGGTTTGTAAC
CTTCCTAATGCTACGGCCCTTTAATACGGTTTCTCACGTTGCGATGACAC
ACACCTCCATCATAATATTATTTTGTTGTTACTTCCTAGCTGTAGTTGTG
CTACTGTTTTGAATCATAATGTAAGTATCTGTGGTTTCTGATGGTCTCAG
GCAACCTCC
```

V$CTCF.04

Rat: BLAT analysis of mouse sequence submitted to rat genome

```
>rn6 dna range=chr2:121078390-121079200
AgAAAAaGTT ATTGTCTTTT CTGTTTTGCT Tttgtttcgc TTTTgACTTg
GGTTGcATTG CTGCACcTTA TTTTATGaCA gGGTCcCgag TAGTcCcGGC
TTTCTgCAAA TTCCCcctTT GTAGcCACGG CTGGCCTTGg ACgTcCCATA
CTCCCTCCga CCTAACCTCC AGAGTGCTtA GgTTACAGGT GTCTGCCACG
cACTgTAACC TTTGaCTGCT TTATTCTtTG CATCtCCTTC CTtctgcgta
tcAGGGTGCC TGCCTTCTCT gCaCCCTTGc AgAGAGCCTG GGTacgCTTT
CCAACCAGTG GAGGCGCTGT TGGCgGCCCC ATGCTcatTC TCcCctcGTT
GACgTTtCAT GGTaAGTGGG tCCTGTTCAG ACCTCTCaAA GCTCTGCTCa
CTTCACTAGc tcccaatcgc cCCGCCTCTG GGgAaCCCCg GtGCCTGCTC
gctgctgttc tgccagggaa aggctgaTTA GCCTCAAGCT CACTCAGGGA
ccTGAGaCCT GGTGCATACA CACTTCTTtC CCAcTaTCTt TCCCAGAAcA
CCCAaTTTGT CTTTAGAAaT gAgAGCTgGA ATGGCAACAg TGAAATCAGC
```

```
TTCAGcccGT AGCTTGgATG CtTTTCATTT TCTGcTTTTG TCTATGCCAG
TGGTTcGTAA CCTTCCTAAT GCTgCGaCCC TTTAATACag ttgttcctcc
tgttgtggtg ACACCcCCAT CATAATATTA TTTTaTcGTT ACTTCCTAaC
TGTAGTTtTG CTACTGTTcT GAATCATAAT GgAAacAaCT GTGGTTTCTG
ATGGTCTCAG
```

V$CTCF.02

#### 2. Sox2-proximal (ubiquitous binding) site (labeled 5, Fig. 4A)

Mouse: sequence extracted from genome browser

```
>mm9_dna range=chr3:34537878-34538786
GTAATCAAGGTGCTGTATGTATGTATGAAAGGGGTGTGTGTGTTTCTTTT
TAAGCAAAAACTAAACGTCTGTGAGTCCCTATAGGTAAATCCAATAATGA
AAATTTTCGGGTTTCCATCTTAAAACTGTTAAATTCTAGTATTGCTCTCC
TACTAAGATCCTTGTGCAGACCGCAGTGACTTAGATGCTGAAGGCTCAC
TTCGATCCCTGTTGGGACTGGGGGAGTGGGGGTGTGCTGTAATTGGGGTG
AAAGAGAGCTTGAATTTAAAAAAGTGAGACAGGGGACATGACTAACCCTC
ATTCTTTATGCAACTGGAATAGAGGGAGATAGAAACGCAAGGGGGCAGGA
TCATCAAGAAATATTACCTGAGAGGGAATACAGAGGCTGACGAGGCTGGG
ATCTGGGAGGGCTGAGAAAAGCTGGCTAGGGCGCTGGGCTACTCTCTGGC
GCCCTCTAATGACAAATCGGACGACATTACCTTCGCCATCCCACGCTCAG
CAAAGGTTTTTTTTTTTTTTTTGTAGCTGCTGAAGCTGTGGTTTTTGTTG
CGCTTGTTTGGTTGGTTGGTTGGTTGATTTGGTTTGTTTTTGCTTTGCTT
TGCTTTGCTCTGCACTTAGTAGAAAAGATAGGTATTTTTCCGTCCAGAAA
CTTAGTACAGGTGATTTTTGGCGCGCGTATTGCCTCCAAGCTGAGGATGG
CTGGCTGTTTGATTCGTGGAGCGAGGCCCTGTGGGGATGAGCAGTGTGGG
GGGAGGGGGGCTTCTCAGCCTGAGAAGTGTGGTTGGGAAGGCCAGTCTGA
GGATCCTGACACCCGCCCATCATCGCAATTCAAGGAGGCTCCTGAGGACC
TCTGTCACCTAGGCGTGCAGTGGCCAGGGGCCTCCCTTTCCTTCTTTCTC
CTCCTCTTC
```

V$CTCF.04
V$CTCF.04
V$CTCF.01

Rat: BLAT analysis of mouse sequence submitted to rat genome

```
>rn6 dna range=chr2:121078390-121079200
GTcATCAAGG TGtTGTATGg atggATGAAA GGGGTGTGTG TaTTTCTcTT  121154019
TAAGtgaaca ctaagaaaca tCcGTGAGTC CCcATAGGTA AATCCAATAA  121154069
TGAAAAaTTT CGGGTTTCCA TCTTAAAACT GTTAAATTCT AGTATTGCTC  121154119
TCCTACTAAG AgCCcTGTGC AGACCGCAGT GACTTggaat gaaaagtacA  121154169
GATaCTGAAG GGCTCACTTC aATCCCcGTT GatACTGGGG Ggcgggggg  121154219
gggggcgcCT GTAATTGGGG TGAgaAAGAG cttgcatttt tttAAAAAGT  121154269
GAGgCAGGaG ACATGAtgAA CCCTCATTCT TTATGCAACT GaAATAGAGG  121154319
GAGATAGAAA CGCAAGGGGG CAGGATtATC AAGAAcTATT ACCTGAGAGG  121154369
GAATACAGAG GCTGACGAGG CTaGGAgCTG GGAGGGCTGt GAAAAGCTGG  121154419
CTAGGGCGgT GGGCTACTCT CTGGCGCCCT CTAATGACAA ATCGaACGAC  121154469
ATTgCCTTCG CCATCCCACG CTCAGgagag ggttttgtcg ctgtggaggc  121154519
tggggggggg gggggttgt tgttgttgct gttgtgttgc TTGTTGCTTG  121154569
TTTGGTTGGc TGGTTGATTT GGTTTGTatt tgTTTTGCTT TtCTTTtCTC  121154619
TGCACTTAGT AGAAgAGATA GGTATTTTTC aGTCCAGAAA CTTAGTACAG  121154669
GTGATTTTTG GCGCGCGTAT TcCtTCCAAG Ccaaggaaag gatGATGGaT  121154719
GGCTGTTTGA TTCGTGGAGC cAGGCCCTGT GGGGATGAGC AGTGcagTGG  121154769
GGaagGGAGG tTTCTCAGCC TGAGAGTGTG tTTGGGAAGa CCAGTgTGAG  121154819
GgtctgtgcA TCCTGACACC CGCCCATCAT CGCcATTCAA GGAGGCTCCT  121154869
GAGGACCTCT GTCACCTAGG CaTGCAGTGa CCAGGGGCCT CCCTcTCCcT  121154919
CTcTCTCCTC CTCTTC
```

V$CTCF.04
V$CTCF.04
V$CTCF.01

### S9. Conservation of Sox2 ‘brain super-enhancer’ sequence

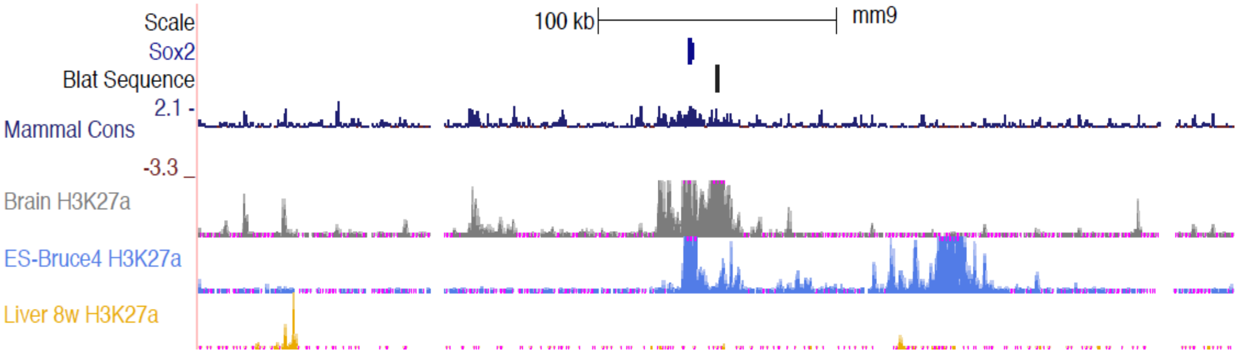


Fig. S9. UCSC genome browser image (mouse, mm9) showing the position of a ChIP target sequence (labeled, Blat sequence) within a Sox2 ‘brain super-enhancer’ (Li et al, 2014) indicated by peaks of H3K27ac association. Note an additional, downstream ES cell-specific super-enhancer (Li et al, 2014). Mammal Cons = Placental mammal basewise conservation (phyloP).

Mouse: BLAT sequence taken from genome browser

```
>mm9_ dna range=chr3:34560628-34561261
CGCAGCTCTCTTCCCTCCTTTCGGGCCTGTGGTGTCGGCCTCTTTTGCTCCGGCTGACGATGCGGGTGG
GTTTGCTGGTGAAAGTGAAGACGTCGCAGTAGCTGTCACCCAGTTCTGCTGTACTTTAAAGCGCTCGCT
ATTGTTTGATTACTAATTCGAGTTAATATGATCTTTATGGCAACATAACAGTGGATAATTGGCCTTTTT
TCTCAACAACAATGTAAATCACAATCGCTTTATTATTTAATAACGTCAAACGCTTCGTTGGATTGGCCC
CGACCGAGAAGACCTTTTCTTTCTAGACAGTATTATGGGGGAAGGACAATGAAGATCTAATTGTGGAGG
TTTCATAAGTTGGCTCCGGTAAATGAACAGCTTCGATATGGAAGTCCCAGCAGATGTGGGCACCGCTAG
CTGTAGAAACTCCACCCCACCCCTCTACCCACCATCCAAACACACTGTTGGAGGCTTTCTGGGGTGGCT
GTGGTCCACGGAGCCTAGCAACCACCTGAAGTTCCCAGTAGCAATTAGA
```

Rat : BLAT analysis of mouse sequence submitted to rat genome reveals a high level of sequence conservation (identical bases in blue).

```
>rn6 dna range=chr2:121176856-121177384
CGCAGCTCTC TTCCCTCCTT TCGGGCCTGT GGTGTCGGCC TCTTTTGCTC  121176906
CGGCTGACGA TGCGGGTGGG TTTGCTGGTG AAAGTGAAGA CGTCGCgGTA  121176956
GCTGTCACCC AGTTCTGCTG TACTTTtAAA GCGCTCGCaT ATTGTTTGAT  121177006
TACaAATTCG AGTTAATATG ATCTTTATGG CAACATAACA GTGGATAATT  121177056
GGCCTTTTTT CTCAACAACA ATGTAAATCA CAATCGCTTT ATTATTTAAT  121177106
AACGTCAAAC GCTTCGTTGG ATTGGCCCCG ACCGAGAAGA CCTTTTCTTT  121177156
CTcGACAGTA TTATGGGGGA AGGACAATGA AGATCTAATT GTGGAGGTTT  121177206
CATAAGTTGG CTCCGGTAAA TGAACAGCTT CGATATGGAA GTCCCAGCAG  121177256
ATGTGcGCAC CGCTAGCTGT AGAAACTCCA CCCCACCCCT CTACCCACCA  121177306
TCCAAACACA CTGTTGGAGG CTTTCTGGGG TGGCTGTGGT CCACaGAGCC  121177356
TAGCAAGAAG TTCCCAGgAG CAATTAGA
```

**S10. Analysis of consensus transcription factor binding sites in Sox2 enhancers with diencephalic activity.**
Consensus sequences present in 3 or more independent locations within rat genomic enhancer sequence are listed:

**U6**
Lhx3, Hand1, S8
**(N3[C4]**
SRY, AP-2alpha, Pax2, Sp1, AP-1, ARP-1, Brn-2
**D1**
Xvent-1
**N2 C(8)**
CdxA, SRY, c-Myb, Sp1, NF-kappaB
**N2 C(9)**
None
**N3 [C2]**
CdxA, STAT5A, Tal-1beta

Sequences were searched using the Lasagna tool (see text). Enhancer nomenclature (U6 etc) is derived from Okamoto et al (2015).

**S11.**

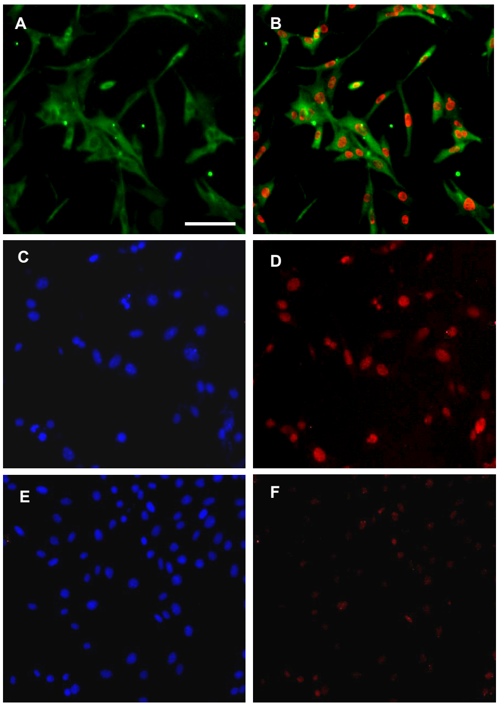


Fig. S11. Over-expression of LHX1 in HT-22 cells. Representative fluorescence microscopic images of fixed, cultured HT-22 cells showing nuclear LHX1 expression following transfection. A&B. Immunodetection of cytoplasmic BetaIII tubulin (green, antibody G7121, Promega), and nuclear histone (H3K27Ac, red, antibody 39133, Active Motif). C-E. Clear immunodetection of LHX1 (red, antibody C6, Santa Cruz) in nuclei of transfected cells (D) compared with a mock transfection (F). Nuclei labeled with DAPI (4',6-diamidino-2-phenylindole) are shown for comparison (C,E). Scale bar: 50µm.

**S12.**

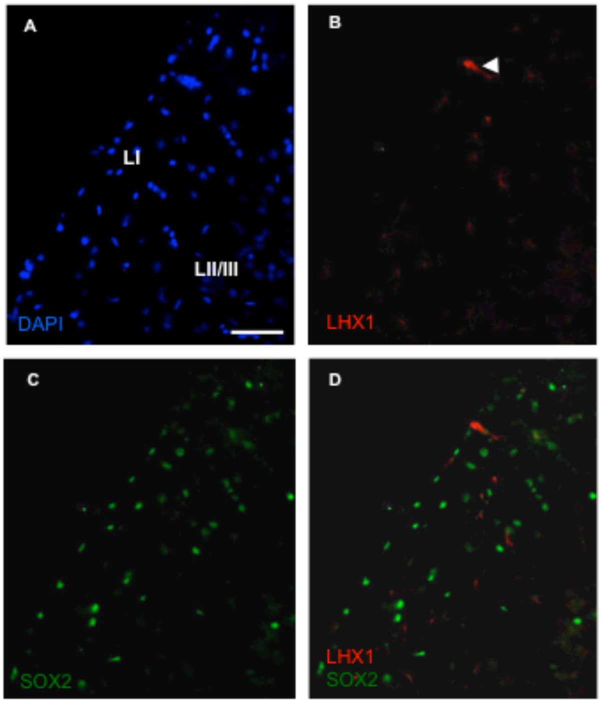


Fig. S12. LHX1 is not expressed in the adult rat brain cortex, but SOX2 is expressed in scattered neurons. Representative fluorescence microscopic images of male PN50 brain showing LHX1 and SOX2 immunoreactivity in neurons. Note the limited detection of blood vessels following the LHX1 immunohistochemical procedure (one example indicated by arrowhead), and a scattered distribution of SOX2+/LHX1-negative neurons. Abbreviations: DAPI, 4',6-diamidino-2-phenylindole; LI, layer I of cortex, LII/LII partial view of cortical layers II and III. Scale bar: 50µm.

| Name | Method | Sequence (5'->3') |
|---|---|---|
| ActbF | PCR | TCATGCCATCCTGCGTCTGGACCT |
| ActbR | PCR | CCGGACTCATCGTACTCCTGCTTG |
| GrinF1 | PCR | TCCACCTGAGTTTCCTTCGC |
| GrinR1 | PCR | GAACCACATCATCCTGCTGG |
| H19F1 | Control ChIP | GATCGTGAAGGGCGCAAGAC |
| H19R1 | Control ChIP | CGTGTGCATCTCTGGAGTGG |
| LhxF1 | PCR | CAACTGGAGACGTTGAAGGC |
| LhxR1 | PCR | GAGATGAGAAAGGTGGACTG |
| SoxF2 | PCR | ACCCGCAGCAAAATGACAGC |
| SoxR2 | PCR | TCCTTCCTTGTCTGTAACGG |
| SoxotF8 | PCR | CCTGCACAGGGCTGGCTGTG |
| SoxotR3 | PCR | ATGGTCCTTCTACTTCTGGG |
| SoxotF1B | PCR | ACAGAGGAGAAGCGGCTGAG |
| SoxHF2 | H3K27ac ChIP | TGAAAGTGAAGACGTCGCGG |
| SoxHR2 | H3K27ac ChIP | ATCCAACGAAGCGTTTGACG |
| SoxCF1 | CTCF ChIP | GGTTACAGGTGTCTGCCACG |
| SoxCR1 | CTCF ChIP | GCTAGTGAAGTGAGCAGAGC |
| SoxCF2 | CTCF ChIP | TTGGCGCGCGTATTCCTTCC |
| SoxCR2 | CTCF ChIP | CTGGTCACTGCATGCCTAGG |
| U6F1 | PCR | CCAGAAGAACAAACGGTATG |
| U6R1 | PCR | GGACTGTGCCAGCAAACATG |
| U6F2 | PCR | CAGGTGCTCATTGCTATAGG |
| U6R2 | PCR | GGACATGTGCCTGTAATTGG |
| U6TFF1 | PCR cloning | ggtaccGTGAAATACTGCCTTCACTG |
| U6TFR3 | PCR cloning | gctagcGCTTCAGTAATTGGCTCAGG |

**Table. S1. Oligonucleotide sequences and use.**
Sequences in lower case indicate introduced restriction enzyme sites.
